# Histopathology and spatial transcriptomics jointly map myofiber-specific pathological programs in mTORC1-driven myopathy

**DOI:** 10.64898/2025.12.03.692123

**Authors:** Jer-En Hsu, Qingyang Zhao, Weiqiu Cheng, Hyun-Min Kang, Susan V. Brooks, Myungjin Kim, Jun Hee Lee

## Abstract

Skeletal muscle is a structurally organized and functionally diverse organ composed of heterogeneous myofiber types and supporting non-myocyte populations that act in concert to generate force, regulate metabolism, and maintain systemic homeostasis. Myopathies occur in many different diseases, but the mechanisms that drive these muscle pathologies are still largely unknown, partly because conventional approaches cannot link histopathological features to molecular states at single-fiber resolution. To address this challenge, we brought histopathology and spatial transcriptomics together by applying high-resolution Seq-Scope technology to a rodent model of mTORC1 hyperactivation, which produces diverse pathological alterations within individual myofibers. Cross-sections from extensor digitorum longus (EDL) and soleus (SOL), two muscles with distinct fiber-type compositions, were profiled to determine how transcriptome changes are linked to histopathological outcomes. Our analyses reveal that mTORC1 hyperactivation elicits distinct, fiber-type–dependent pathological programs. Type I and IIa fibers, abundant in SOL but scarce in EDL, were largely resistant to mTORC1-induced pathology, exhibiting only minimal morphological alterations and no fiber type-specific responses beyond those commonly observed throughout the tissue. In contrast, type IIx fibers, shared between both muscles, diverged into opposing fates: in SOL, they underwent abnormal enlargement driven by sustained growth signaling, cytoskeletal remodeling, and impaired proteostasis with defective autophagy; whereas in EDL, they developed basophilia characterized by lipid-supported respiration fueling excessive ribonucleotide synthesis and RNA accumulation. Within the same muscle, type IIb fibers displayed striking heterogeneity with discrete transcriptional states encompassing canonical stress responses, oxidative metabolic activation, and developmental reprogramming. In parallel, non-myocytic populations, including activated macrophages and fibroblasts, accumulated preferentially in SOL, forming a fibrotic microenvironment supporting inflammation, tissue remodeling and hypertrophy. Taken together, these findings reveal that sustained mTORC1 signaling disrupts muscle homeostasis through distinct metabolic and structural routes, directly linking histopathological phenotypes to their molecular states at single-fiber resolution.

## Introduction

Skeletal muscle is a multifunctional organ that drives movement while being critical for maintaining metabolic homeostasis. This balanced control of essential functions is critical for maintaining health and relies on the heterogeneity of myofibers with respect to contraction and energy metabolism. Differences between myofibers underlie the classic classification into slow and fast, and oxidative and glycolytic fiber types, a framework that has long explained variation in endurance, contractile speed, and fatigue resistance. (1). At the molecular level, this classification is largely determined by the expression of distinct myosin heavy chain (MyHC) isoforms, which define the major fiber types: type I (MF1; slow oxidative), type IIa (MF2A; fast oxidative), type IIx (MF2X; intermediate fast glycolytic), and type IIb (MF2B; fast glycolytic). The functional specialization of each fiber type allows skeletal muscle as a whole to support diverse physiological demands (2).

Mammalian target of rapamycin complex 1 (mTORC1) is a central regulator of cell growth and anabolism that senses inputs from nutrient, growth factor, and energy signals (3). Its kinase activity is activated at the lysosomal surface through the small GTPase RHEB, which is inhibited by the TSC1/TSC2 complex during growth factor deprivation or energy depletion. In parallel, mTORC1 recruitment to lysosomes is controlled by amino acid availability. Under amino acid insufficiency, the GATOR1 complex (DEPDC5, NPRL2, and NPRL3) maintains the Rag GTPases (RRAGA/B and RRAGC/D) in their inactive GDP-bound state, thereby preventing mTORC1 from translocating to the lysosomal surface and becoming activated (4,5). Disruption of either TSC1 or DEPDC5 elevates mTORC1 activity, but full constitutive activation requires disruption of both pathways due to the presence of negative feedback mechanisms (6–8). Once active, mTORC1 phosphorylates downstream targets such as S6K1 and 4E-BP1 to promote mRNA translation, while simultaneously suppresses autophagy initiation through inhibitory phosphorylation of ULK1 (9–11). By linking environmental inputs to biosynthetic outputs, mTORC1 ensures that cell growth and anabolic metabolism occur only under permissive conditions.

In skeletal muscle, appropriate mTORC1 activity is indispensable for growth, maintenance, and repair (12). Muscle-specific deletion of Raptor, an essential subunit of mTORC1, results in profound atrophy and impaired force generation (13), highlighting the importance of proper mTORC1 activity in maintaining muscle homeostasis. On the contrary, sustained hyperactivation of mTORC1, whether induced experimentally through muscle-specific TSC1 deletion or arising naturally during aging, leads to progressive myopathy marked by fiber degeneration, metabolic stress, impaired proteostasis, and loss of regenerative capacity (8). In our previous work, we produced a mouse strain with depletion of both TSC1 and DEPDC5 in muscle tissue (*Ckm-Cre Tsc1^F/F^ Depdc5^F/F^*) to enforce constitutive hyperactivated mTORC1 signaling pathway (6,14). Unlike single-knockout models (7,8), which showed little pathology in young mice, *Ckm-Cre Tsc1^F/F^ Depdc5^F/F^* mice develop severe early-onset myopathy and skeletal muscle dysfunction. This phenotype was associated with excessive oxidative stress, impaired proteostasis, and accumulation of the autophagy adaptor SQSTM1/p62, resulting in dysregulation of myofiber homeostasis.

With recent advances in technologies, single-cell (sc) and single-nucleus (sn) RNA-seq resources have mapped aging muscle across lifespan and diseases, providing valuable insights into cellular composition and aging-associated shifts in regenerative, immune, and fibrotic states (15–26). Spatial transcriptomic and imaging-based methods such as MERFISH (27) and smFISH (28) have further visualized the distribution of specific transcripts within intact myofibers and tissue sections, while multiplex proteomic platforms like CODEX (29) have profiled fiber-type mosaics at the protein level. Together, these approaches have greatly expanded our understanding of muscle biology and aging. However, they fall short in resolving the fiber-type complexity of myopathy. Sc/snRNA-seq relies on tissue dissociation and cannot resolve intact myofibers, while spatial methods to date have been constrained to targeted panels or low resolution, making it difficult to capture the full complexity of fiber-type heterogeneity in the myopathic process.

To directly address this gap, we applied Seq-Scope, a high-resolution spatial transcriptome profiling technology that we recently developed (30,31). Its submicrometer-resolution with unbiased whole-transcriptome coverage enable high-fidelity mapping of transcriptional diversity at the single-myofiber level. Using Seq-Scope, we profiled cross sections of extensor digitorum longus (EDL) muscles and soleus (SOL) muscles, two anatomically distinct tissues encompassing the major myofiber types, from *Ckm-Cre Tsc1^F/F^ Depdc5^F/F^* and control littermates. Our analyses revealed that mTORC1 hyperactivation elicits both global stress responses and fiber-type-specific remodeling programs. Comparison between EDL and SOL demonstrated striking context-dependent plasticity of MF2X, which engaged distinct metabolic and cytoskeletal remodeling pathways in each muscle. Within the same muscle, MF2B reprogrammed their classical glycolytic transcriptome, acquiring either MF2X-like oxidative signatures or undifferentiated myofiber-like developmental signatures. In addition, classical myopathic features, such as abnormal myofiber enlargement and basophilic fibers, were each linked to characteristic transcriptomic profiles with predicted functional outcomes. Overall, this study provides a histopathology-guided, single-fiber spatial framework for dissecting fiber-type–specific remodeling under mTORC1 hyperactivation, providing molecular insight into the developmental processes of myopathies and a foundation for future mechanistic studies of muscle physiology and disease.

## Results

### Seq-Scope enables combined analysis of histopathology and single-myofiber transcriptomics

We have formerly generated a mouse model of unregulated mTORC1 hyperactivation by concomitantly deleting *Tsc1* and *Depdc5*, thereby achieving maximal activation of both Rheb and Rag arms of mTORC1 signaling (Figure 1A). mTORC1 hyperactivation in skeletal muscle severely disrupted tissue homeostasis, leading to loss of muscle function associated with various pathologies, including basophilic fibers, enlarged fibers, and centrally nucleated fibers (Figure 1A; white, black and green arrows, respectively). To capture the molecular changes underlying these phenotypes at the resolution of individual myofibers, we used Seq-Scope, a spatial transcriptomics platform with sub-micrometer resolution (average 0.5–0.7 μm), which we have recently developed and optimized (30,31). By aggregating transcripts according to the histology-based myofiber segmentation, spatial transcriptome could be precisely assigned into individual myofibers. This approach preserved fiber morphology and spatial context while revealing and mapping transcriptomic diversity.

**Figure 1.**
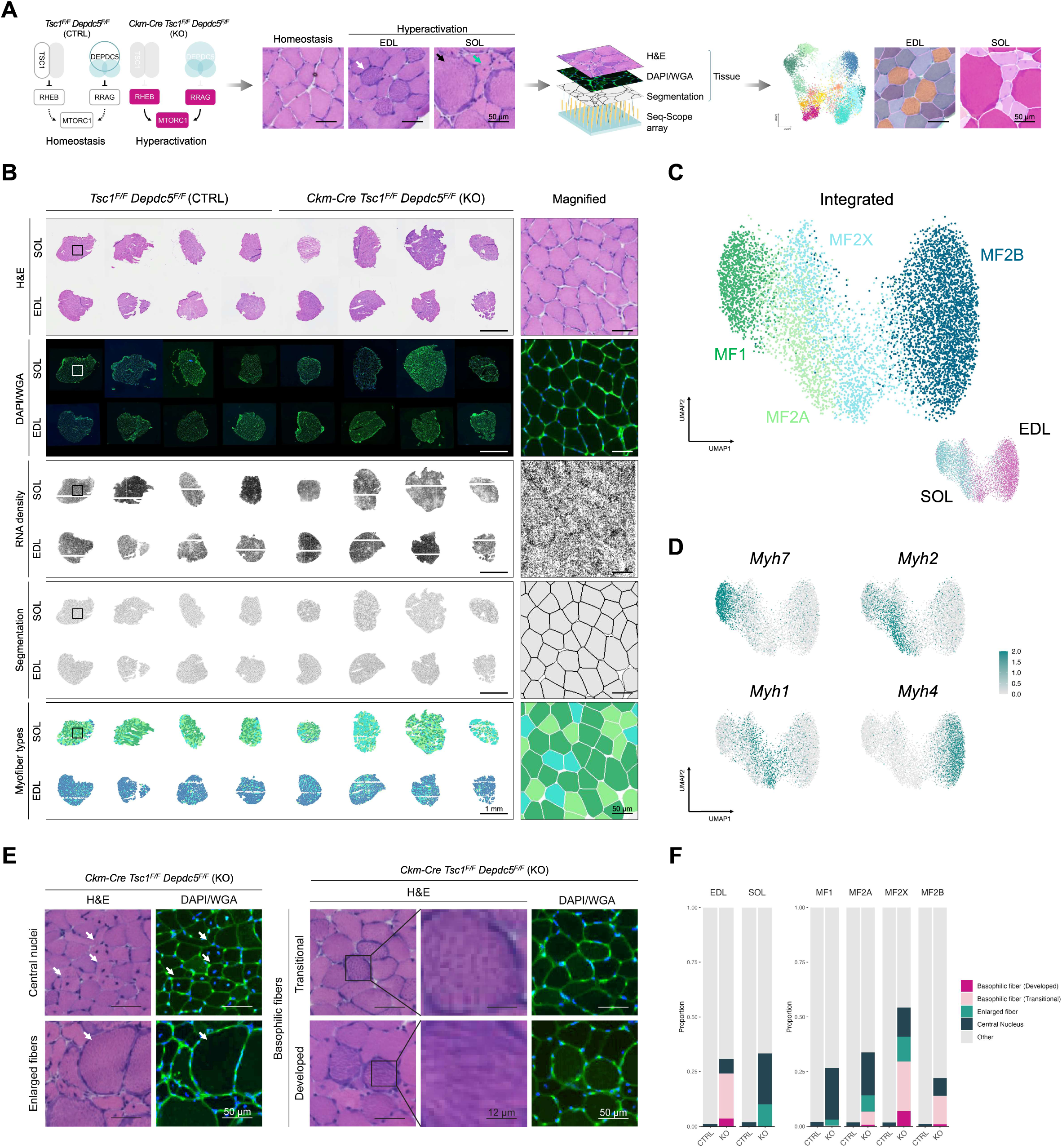
Seq-Scope enables histopathology-guided single-myofiber transcriptome analysis. (A) Schematic of the experimental workflow. Arrows indicate pathological phenotypes including basophilia (white), hypertrophy (black), and central nuclei (green). Hematoxylin and eosin (H&E) histology, DAPI/wheat germ agglutinin (WGA) imaging, and Seq-Scope whole transcriptome analysis are performed on the same slide, which allows precise segmentation and single-myofiber transcriptome analysis. (B) Cross-sections of control and mutant skeletal muscles examined by Seq-Scope. From top to bottom, images are displayed as H&E histology, DAPI/WGA staining, RNA density map from Seq-Scope transcriptome data, single-myofiber segmentation, and spatial myofiber type projection, according to the clusters identified in (C). *Tsc1^F/F^ Depdc5^F/F^*: control; *Ckm-Cre Tsc1^F/F^ Depdc5^F/F^*: mutant. SOL: soleus muscle; EDL: extensor digitorum longus muscle. MF1: myofiber type I; MF2A: myofiber type IIa; MF2X: myofiber type IIx; MF2B: myofiber type IIb. (C) UMAP manifolds displaying the major myofiber types identified by multidimensional clustering. Datasets were integrated across mice. Each point (n = 8616) represents an individual myofiber and is colored according to its corresponding myofiber type (center) or its originating muscle depot (lower right). (D) UMAP manifold colored by the expression levels of myosin heavy chain (Myh) isoforms that represent each myofiber type. (E) Pathological phenotypes observed in mTORC1 hyperactivated skeletal muscle. Left panel: centrally nucleated myofibers (arrows in the top row) and hypertrophic fibers (arrows in the bottom row). Right panel: transitional basophilic fibers (top row, scattered dark particles) and developed basophilic fibers (bottom row, dense and strip-like filling). Boxed areas are magnified on the right. (F) Quantification of pathological phenotype prevalence by muscle depot (left) or by myofiber type (right). “Other” includes myofibers from control muscles as well as morphologically normal fibers from mutant muscles. CTRL, control; KO, mutant.

In this study, four *Ckm-Cre Tsc1^F/F^ Depdc5^F/F^* (muscle-specific deletion of *Tsc1* and *Depdc5*) and four *Tsc1^F/F^ Depdc5^F/F^* (Cre-negative control) littermates were sacrificed at 10 weeks of age. EDL and SOL muscles were harvested, embedded in OCT, freshly frozen, and sectioned. A total of 16 cross-sections (two muscles per mouse) were placed onto Seq-Scope arrays for spatial transcriptomic profiling. The exact same sections were subjected to H&E and DAPI/WGA imaging on the array surface prior to transcript capture and library generation (Figure 1B; first and second row, respectively). Sequencing of the Seq-Scope library yielded a total of 700 million unique reads that successfully aligned to the mm10 reference genome with valid spatial barcodes. The resulting RNA density map (Figure 1B, third row) revealed strong localization of transcripts at the periphery of myofibers, consistent with known muscle architecture. Using WGA imaging data and Cellpose segmentation algorithm to delineate cellular membranes, we segmented 8,616 individual myofibers across 16 tissue sections (Figure 1B, fourth row, S1A). Data from each sample was considered independent, integrated and clustered to assign myofiber identities. The data showed minimal batch effects, aside from the expected separation of SOL and EDL with their distinct myofiber compositions (Figure 1C, S1B–D). Myofiber types were also clearly distinguished by the expression of their corresponding myosin heavy chain isoforms (Figure 1D, S1C) and by metabolic genes associated with oxidative (MF1/MF2A) or glycolytic (MF2B) pathways (Figure S1E–F).

To systematically assess the histopathological features observed in mutant muscle, we quantified the prevalence of basophilic fibers, enlarged fibers, and centrally nucleated fibers across all sections (Figure 1E). Basophilic fibers were further classified by staining patterns: transitional basophilic fibers showed scattered dark particles (Figure 1E, right panel, top row), whereas developed basophilic fibers displayed dense, strip-like filling (Figure 1E, right panel, bottom row). Strikingly, basophilic fibers were observed only in mutant EDL muscle, while enlarged fibers were restricted to mutant SOL muscle (Figure 1F, left panel). Fibers with central nuclei were also more frequent in mutant SOL muscle. When stratified by fiber type, we found that MF2X fibers showed the highest frequency of pathological features relative to other fiber types, suggesting that MF2X fibers exhibit the strongest pathological response to mTORC1 hyperactivation (Figure 1F, right panel, S1G).

### Identification of fiber type-specific responses to mTORC1 hyperactivation

Skeletal muscle fiber types differ not only in contractile speed but also in metabolic specialization (1). These intrinsic differences suggest that individual fiber types may respond unevenly to stress. To test this, we conducted differential gene expression analysis to compare transcriptomic signatures of each myofiber cluster between control (homeostasis) and mutant (hyperactivation) muscle (Figure 2A, S2A). Even though many genes were commonly upregulated in response to mTORC1 hyperactivation, representing conserved response pathways, substantial number of genes showed myofiber type–specific signatures (Figure 2B). As described in the Methods, we have classified the genes upregulated under mTORC1 hyperactivation into global response (expressed across all four fiber types, 25%); partially specific response (expressed in two or three fiber types, 49.2% in sum); and responses specific for MF2A (5.6%, 7 genes), MF2X (4.0%, 5 genes), and MF2B (16.1%, 20 genes) fiber types (Figure 2C, S2B-C). Notably, MF1 transcriptome only showed global responses, without any unique signatures, consistent with its well-documented resilience to metabolic and structural stress (32).

**Figure 2.**
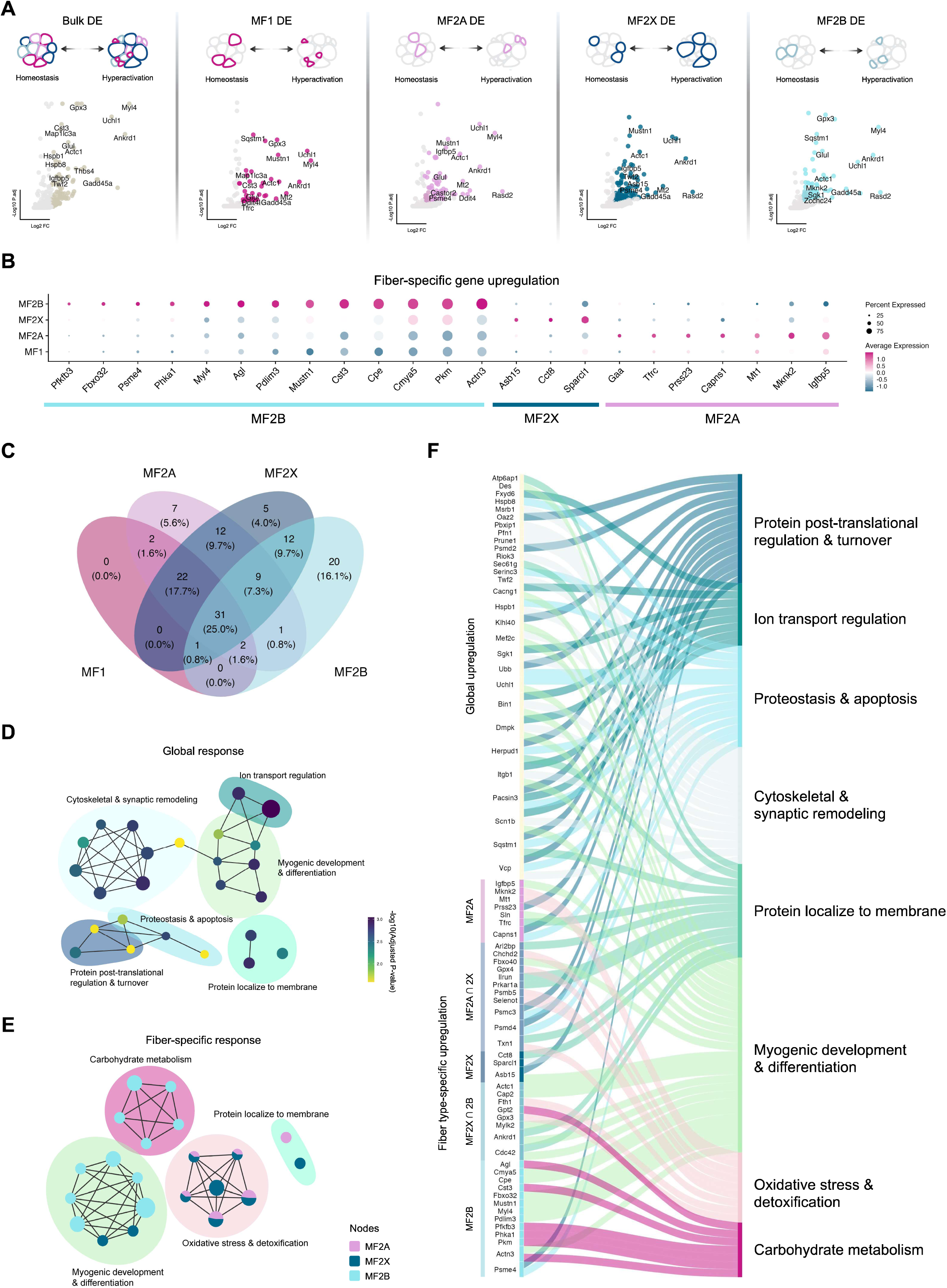
Identification of fiber type-specific responses to mTORC1 hyperactivation. (A) Schematic of the analytical workflow for fiber type-specific differential expression (DE) analysis comparing control (homeostasis) and mutant (hyperactivation) muscle. Upregulated genes are shown (adjusted p-value < 0.05, log2FC > 1.5). (B) Representative expression patterns of genes that are exclusively upregulated in individual myofiber types, selected from a broader set (Fig. S2B). Exclusivity was determined by the Kruskal-Wallis test (adjusted p-value < 0.05). Exclusively expressed genes were not identified in mutant MF1 fibers. (C) Venn diagram depicting the uniqueness and overlap of upregulated genes across different myofiber types, with numbers and percentages shown for both shared and unique gene sets. (D) Genes upregulated across all four fiber types (Global Response) were analyzed by gene ontology (GO) enrichment and visualized as a network. Each node represents a GO term, grouped by functional proximity within the enrichment hierarchy. Node size indicates the number of genes associated with the term, and node color reflects enrichment significance (adjusted p-value). (E) Genes selectively upregulated in specific fiber types were analyzed using compareGO. GO terms were grouped by functional proximity. Node size reflects the number of contributing genes, and node colors are shown as pie charts indicating the relative enrichment strength (adjusted p-value) contributed by each fiber type. (F) Genes globally upregulated across all fiber types and genes selectively upregulated in specific fiber types were mapped to enriched GO groups. The origin of each gene set is indicated by the stratum, and connecting threads represent gene-to-pathway relationships, colored by GO term identity.

To systematically evaluate the functional implications of both global and fiber-type–specific signatures, we performed gene ontology (GO) enrichment analysis. Globally expressed genes were enriched in pathways related to ion transport regulation, muscle differentiation, cytoskeletal and synaptic remodeling, protein localization to membranes, and proteostasis (Figure 2D). These pathways, directly linked to mTORC1 hyperactivation-induced proteostasis disruption and damage responses, were also observed in our previous bulk RNA-seq findings (6), highlighting the reproducibility of our analysis across different platforms.

In contrast, enabled by the Seq-Scope technology, the present study revealed novel findings of fiber type–specific responses (Figure 2B, 2E). CompareGO clustering showed that MF2B myofibers, despite experiencing the same mTORC1-driven stress as other fiber types, exhibited little enrichment of oxidative stress or detoxification processes. Instead, their response was dominated by reinforcement of carbohydrate metabolic pathways with the upregulation of *Pkm*, *Phka1*, *Agl*, and *Pfkb3*, indicating a rigid glycolytic program even under pathological conditions. By contrast, MF2A myofibers enriched heavily in stress-response and protein quality-control pathways, reflected by the induction of *Tfrc* (transferrin receptor, iron uptake), *Mt1* (metallothionein, oxidative stress buffering), *Psmd4*, and *Psmc3* (proteasome-related subunits), alongside regulators such as *Igfbp5* (insulin growth factor binding protein) and *Sln* (sarcolipin, calcium handling) that mark contractile and developmental specialization. MF2X myofibers, on the other hand, exhibited strong remodeling signatures with elevated *Asb15* (E3 ubiquitin ligase adaptor in myogenesis), *Sparcl1* (extracellular matrix glycoprotein), and *Actc1* (cardiac α-actin, muscle development marker), accompanied by stress regulators *Ctsb* (cathepsin B) and *Gpx3* (glutathione peroxidase), suggesting greater transcriptional plasticity in structural and oxidative adaptation. Together, these results establish a clear correspondence between gene-level signatures and higher-order functional pathways (Figure 2F), validating the concept that different fiber types deploy distinct transcriptional programs in response to mTORC1 hyperactivation.

### Distinct mTORC1-driven adaptations of type IIx myofibers in different muscles

Myofibers type IIx (MF2X), known to express *Myh1* isoform, are often viewed as an intermediate fast-twitch population positioned between oxidative MF2A and glycolytic MF2B with respect to contractile and metabolic features (1). Importantly, MF2X represents the only fiber type consistently present in both the predominantly oxidative SOL muscle and the glycolytic EDL muscle, positioning it as a shared node across otherwise distinct fiber-type compositions (33). Recent snRNA-seq studies in both mouse and human muscle have confirmed that MF2X nuclei form a distinct transcriptional cluster, separable from myofiber MF2A and MF2B, and that their representation and state can shift under different physiological and pathological contexts such as regeneration or denervation (17,28). These datasets highlight that MF2X are not a homogeneous class but instead display notable context-specific divergence.

To investigate how such a dynamic fiber type responds to mTORC1 hyperactivation, we isolated total of 1,447 MF2X myofibers from the dataset, representing all different tissue samples that integrated well with each other (Figure S1B, 3A, and S3A). Although control MF2X fibers from SOL and EDL are mixed well in the UMAP manifold (Figure 3A, left), mTORC1-hyperactivated MF2X fibers showed two distinct clusters, representing damage-associated MF2X-SOL and MF2X-EDL fibers (Figure 3A, right). Notably, MF2X-SOL was predominantly associated with myofiber enlargement (Figure 3B, top panel) while MF2X-EDL often exhibited basophilia (Figure 3C, bottom panel).

**Figure 3.**
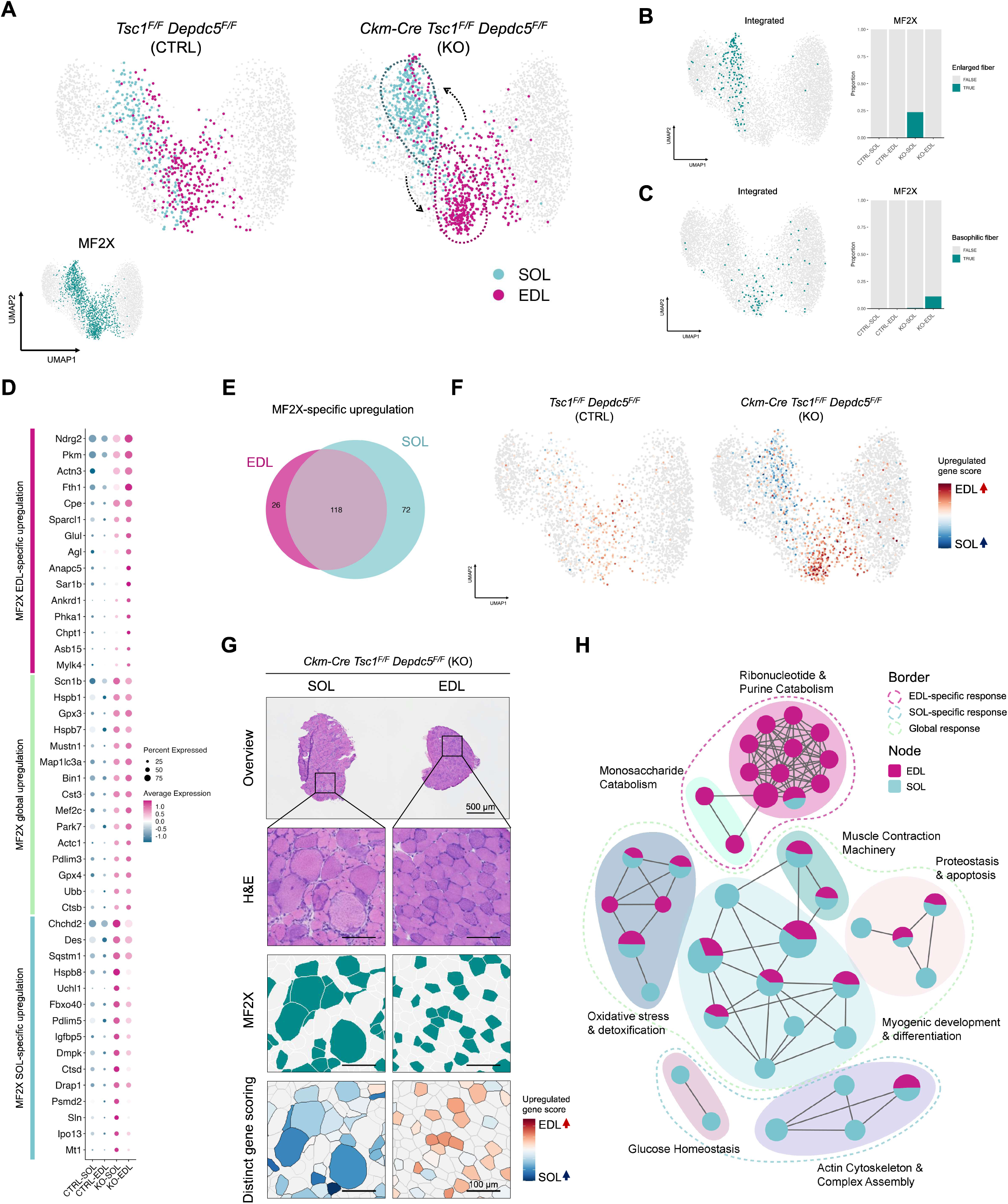
Distinct hyperactive mTORC1 responses of type IIx myofibers in different muscles. (A) UMAP visualization of myofiber datapoints separated as control (left) or mutant (right) groups, with muscle identity (EDL: red, or SOL: blue) annotated within the original MF2X depot (lower left). All other fiber types were greyed out. Dotted arrows and shapes indicate the bidirectional shift of the MF2X subclusters in response to mTORC1 hyperactivation. (B-C) Enlarged fibers (B) and developed basophilic fibers (C) were marked in the UMAP manifold (left). Their proportions were quantified from MF2X cluster of indicated muscles, and presented in a bar graph (right). (D) Representative expression patterns of genes that are upregulated in response to mTORC1 hyperactivation in all MF2X fibers (MF2X global upregulation) or specifically in MF2X of EDL or MF2X of SOL, selected from a broader set (Fig. S3C). Exclusivity was determined using pairwise Wilcoxon tests (adjusted p-value < 0.05). (E) Venn diagram depicting the uniqueness and overlap of genes upregulated in MF2X of SOL and MF2X of EDL. (F) Cumulative module score of muscle-specific upregulated genes displayed in integrated UMAP manifold with control (left) and mutant (right) myofiber datapoints separated. (G) Spatial images of representative mutant sections from each muscle. Boxed areas are magnified below (H&E). MF2X myofiber identity (MF2X) and cumulative module scores (Distinct gene scoring) representing the axis of SOL-and EDL-specific responses are also visualized at the myofiber level. (H) GO enrichment network representing group of genes that are upregulated in response to mTORC1 hyperactivation in all MF2X fibers (Global response) or specifically in MF2X of EDL (EDL-specific) or MF2X of SOL (SOL-specific). Pathways were grouped by functional similarity. Node size reflects the number of genes associated with each pathway, and node colors are shown as pie charts indicating the relative enrichment significance (adjusted p-values) contributed by each muscle-specific gene set. For the complete results with GO term names, see Fig. S3F.

Differential expression analysis between control and mTORC1-hyperactivated MF2X-SOL or MF2X-EDL revealed fiber type-specific transcriptional responses (Figure S3B). Among 216 genes highly upregulated by hyperactive mTORC1, 26 genes were classified to be EDL-specific, while 72 genes were classified to be SOL-specific (Figure 3D-E, S3C). Other genes were similarly upregulated in MF2X from both muscles. Gene score analysis demonstrated that mTORC1 hyperactivation was the predominant driver of these transcriptional changes (Figure 3F, right), while such differences were minimal in control muscle (Figure 3F, left). Histology and spatial profiling of the MF2X myofibers further validated the distinct transcriptional and morphological separation between mutant SOL and EDL muscles (Figure 3G, S3D). While MF2X-EDL fibers remained small or normal in size, MF2X-SOL fibers frequently exhibited noticeable hypertrophy (Figure 3G, S3E).

Using the SOL- and EDL-specific mTORC1-responsive genes (Fig. 3D, 3E and S3C), we performed gene set enrichment analysis (Figure 3H, S3F). Three categories of pathways emerged: those enriched in global MF2X response (proteostasis & apoptosis, muscle contraction, myogenic development, and oxidative stress response), SOL-specific MF2X response (cytoskeletal complex assembly, and glucose homeostasis), and EDL-specific MF2X response (ribonucleotide/purine metabolism, and monosaccharide catabolism). Together, these findings demonstrate that, even within a shared fiber class, mTORC1 hyperactivation elicits divergent muscle-specific adaptations, underscoring the context-dependent plasticity of type IIx myofibers.

### Type IIx myofibers in Soleus undergo hypertrophy with structural remodeling

Abnormal myofiber enlargement is a pathological hallmark observed in diverse muscle disorders and experimental models (1). As mTORC1 hyperactivation induced myofiber enlargement specifically in SOL muscle (Figure 1F), we investigate its molecular basis by re-analyzing 2,972 SOL myofibers, with a focus on the difference between control and mTORC1-hyperactivated muscle tissues (Figure 4A). In mutant SOL muscles, enlarged fibers were mainly recognized as MF2X, with only a small contribution from MF2A (Figure 4A-B). Fiber size distributions derived from cross-sectional segmentation confirmed that while most MF2X fibers in mutant muscle expanded several-fold compared with controls, MF1 and the majority of MF2A shrank instead (Figure 4C, S4A). These observations are consistent with IHC validation (Figure S3E) and previously reported immunostaining quantification (8). We further noted a universal increase in *Myh1* expression and decrease in *Myh7/2* expression (Figure S4B). The proportion of MF2X fibers in general were elevated in mutant SOL muscle (Figure S4C). Importantly, this shift was observed consistently across independent batches, suggesting a potential fiber-type shift during stress.

**Figure 4.**
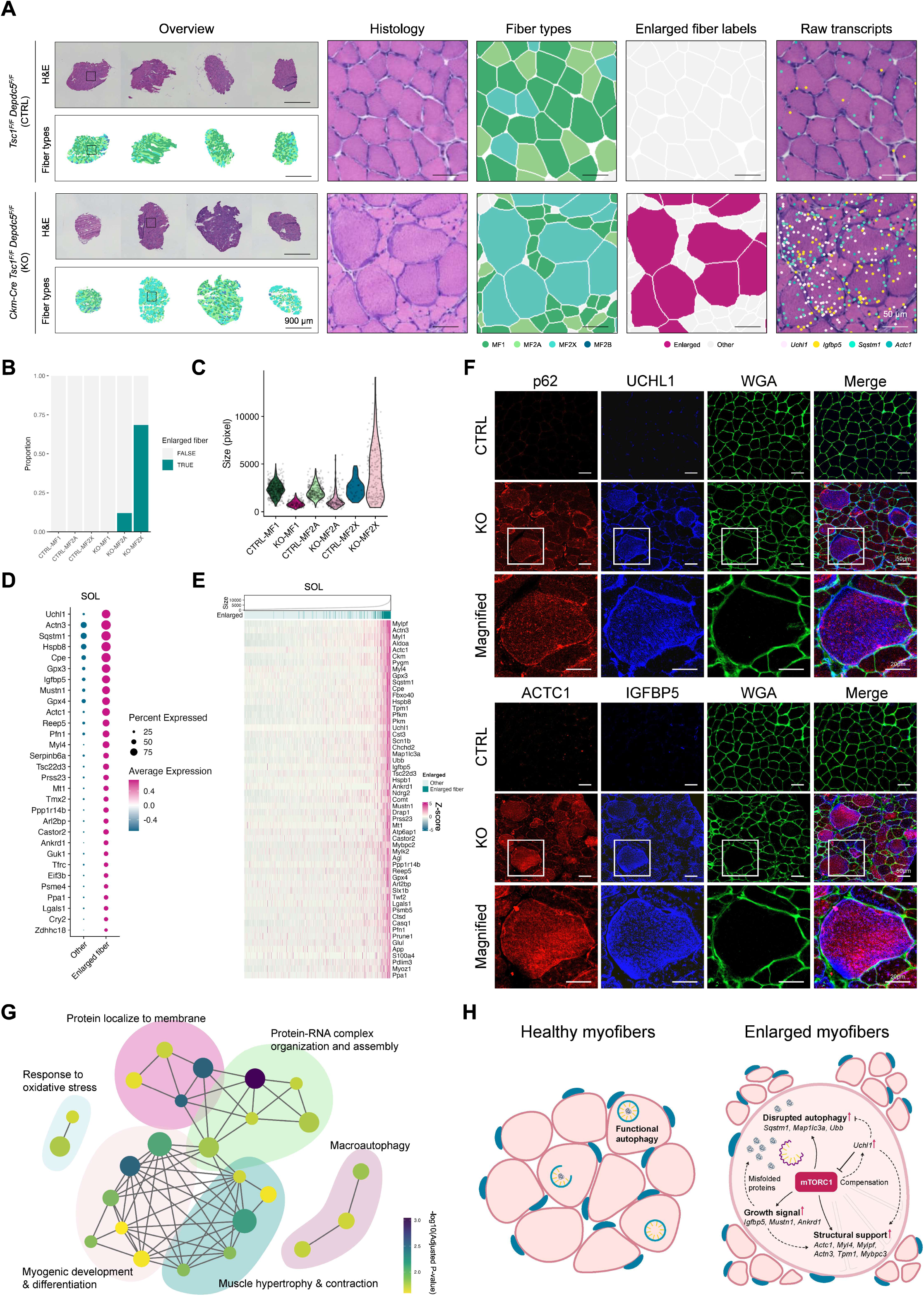
Type IIx myofibers in Soleus undergo hypertrophy with structural remodeling. (A) Spatial images of SOL sections. Left: histology overview of 4 control sections (top) and 4 mutant sections (bottom) overlaid with fiber type images. Right: Spatial images of magnified boxed area from corresponding overview, presenting from left to right as histology, fiber types, enlarged fiber labels, and raw transcripts plotted as colored dots. (B) Prevalence of enlarged fibers in different myofiber clusters of SOL was quantified and presented as a bar graph. (C) Size of fibers in different myofiber clusters of SOL was calculated and presented as a violin plot. One pixel corresponds to ∼0.57 µm^2^. (D) Expression of genes highly upregulated in enlarged fibers were presented in a dot plot. (E) Correlation between fiber size and gene expression. Myofibers (x-axis) are ordered from small to large. Positively correlated genes (y-axis) are ranked by correlation strength. The expression level of the genes is colored based on z-scores. RNA counts within single fiber were normalized by fiber size to estimate RNA density. (F) Immunostaining of indicated proteins in control or mutant SOL muscle. WGA staining (green) outlines myofiber borders. Staining was performed on two serial sections (top and bottom). Boxed areas highlighting an enlarged fiber are magnified at the bottom. (G) GO enrichment network based on genes upregulated in enlarged fibers. Pathways were grouped by functional similarity. Node size corresponds to the number of genes associated with each pathway, and node color reflects enrichment significance (adjusted p-value). For the complete results with GO term names, see Fig. S4F. (H) Schematic diagram illustrating transcriptome regulation underlying pathological hypertrophy in mTORC1-activated type IIx myofibers in SOL.

In addition to exhibiting prominent hypertrophy, MF2X fibers also displayed more pronounced changes at the molecular level. Among the genes differentially expressed between control and mutant SOL tissues, MF2X fibers showed the strongest overall expression changes compared with other fiber types (Figure S4D). Since MF2X fibers showed both abnormal enlargement and transcriptome alterations, we further evaluated the molecular basis underlying the mTORC1-driven enlarged fibers, which occurs only in a subset of MF2X. To address this, we directly compared enlarged and non-enlarged fibers, enabling clearer identification of genes associated with mTORC1-driven hypertrophy (Figure 4D). These genes were generally expressed at low levels in non-hypertrophic fibers and their expressions showed strong positive correlations with myofiber sizes (Figure 4E), even after normalization to each myofiber’s RNA content (Figure S4E).

Genes strongly correlated with fiber enlargement included classical autophagy regulators such as *Sqstm1* (p62), *Map1lc3a* (LC3A) and *Ubb*, as well as proteasome or protease components such as *Psmb5* and *Ctsd,* suggesting an association between myofiber hypertrophy and mTORC1 hyperactivation-driven proteostasis disruption. Genes involved in muscle growth and regeneration (e.g., *Igfbp5*, *Mustn1* and *Ankrd1*) and structural genes indicating sarcomeric and cytoskeletal remodeling (e.g., *Actc1*, *Myl4*, *Mylpf*, *Actn3*, *Tpm1* and *Mybpc3*) were also specifically upregulated in hypertrophic myofibers from mTORC1-hyperactivated muscles.

In addition, *Uchl1*, a ubiquitin C-terminal hydrolase, was strongly correlated with fiber enlargement. Notably, Uchl1 has been shown to act as a brake on mTORC1 signaling in skeletal muscle, where its deletion results in sustained mTORC1 activation, impaired autophagy, and progressive muscle atrophy (34). Consistent with this, UCHL1 is also required for maintaining ubiquitin homeostasis and proper differentiation in myotubes, with knockdown leading to protein aggregation (35). Its elevated expression in enlarged fibers therefore suggests a compensatory role in managing proteostasis and restraining excessive mTORC1 activity under hyperactivation.

Importantly, the specific upregulation of *p62*, *Igfbp5*, *Actc1*, and *Uchl1* in hypertrophic myofibers was confirmed through IHC staining of their protein products (Figure 4F). This coordinated program, combining impaired proteostasis, persistent growth signaling, and cytoskeletal remodeling, was reflected in our GO network analysis (Figure 4G, S4F). Together, these results connect the molecular changes to the enlargement myopathy observed in SOL muscle (Figure 4H).

### Central nuclei are prevalent across various types of myofibers

The presence of centrally located nuclei in skeletal myofibers is a well-recognized histopathological feature, commonly associated with regeneration, myopathies, and various muscle stress conditions (36,37). Histological analysis revealed a higher proportion of centrally nucleated myofibers in mutant SOL compared with mutant EDL muscles (Figure 1F, S4H, left panel). Although no strong fiber-type specificity was detected, oxidative fibers, such as MF1 and MF2A, had a subtle tendency to show more frequent central nuclei compared to the glycolytic fibers (Figure S1G, S4G, and S4H, right panel). To assess whether central nuclei regions display distinct transcriptional features, we segmented 430 central nuclei from 2 mutant EDL and 2 mutant SOL muscles using combined H&E, DAPI, and WGA staining. Nuclei were dilated by 2, 4, or 6 µm to capture sufficient transcripts, and their transcriptomes were compared with 24-µm hexagon profiles from the 4 selected sections to generate a spatial single-nuclei–like dataset (Figure S4I-J). The analysis did not reveal strong enrichment for a specific gene or pathway, likely reflecting that the central nuclei feature could happen for a variety of different myofibers without representing a specific pathway. However, we were able to identify several zinc finger proteins, such as *Zfp93* and *Zfp429*, which were significantly enriched in central nucleic regions and may be involved in positioning the nucleus during the injury (Figure S4K).

### Basophilic fibers activate lipid and oxidative metabolism to sustain elevated RNA synthesis

Basophilic fibers represent a distinctive histological phenotype, defined by enhanced hematoxylin accumulation in the cytoplasm, reflecting elevated nucleotide content and often emerging under stress conditions (38). Here, based on H&E histology, we identified and isolated 48 developed basophilic fibers and 387 transitional basophilic fibers where the majority were restricted to MF2X fibers in mutant EDL muscle (Figure 5A-C, S5A, left panel). To validate the notion that more hematoxylin staining reflects elevated nucleotide content, we quantified RNA abundance in single fibers by counting unique transcripts from the spatial transcriptome dataset. Raw counts showed only modest increases in some developed basophilic fibers (Figure S5B) likely due to the presence of hypertrophic fibers, which occupied more fiber volumes in histology, in non-basophilic fiber group. After normalizing RNA counts by fiber area to measure RNA concentration, we then observed a robust elevation of RNA content across both transitional and developed basophilic fibers (Figure 5D-E).

**Figure 5.**
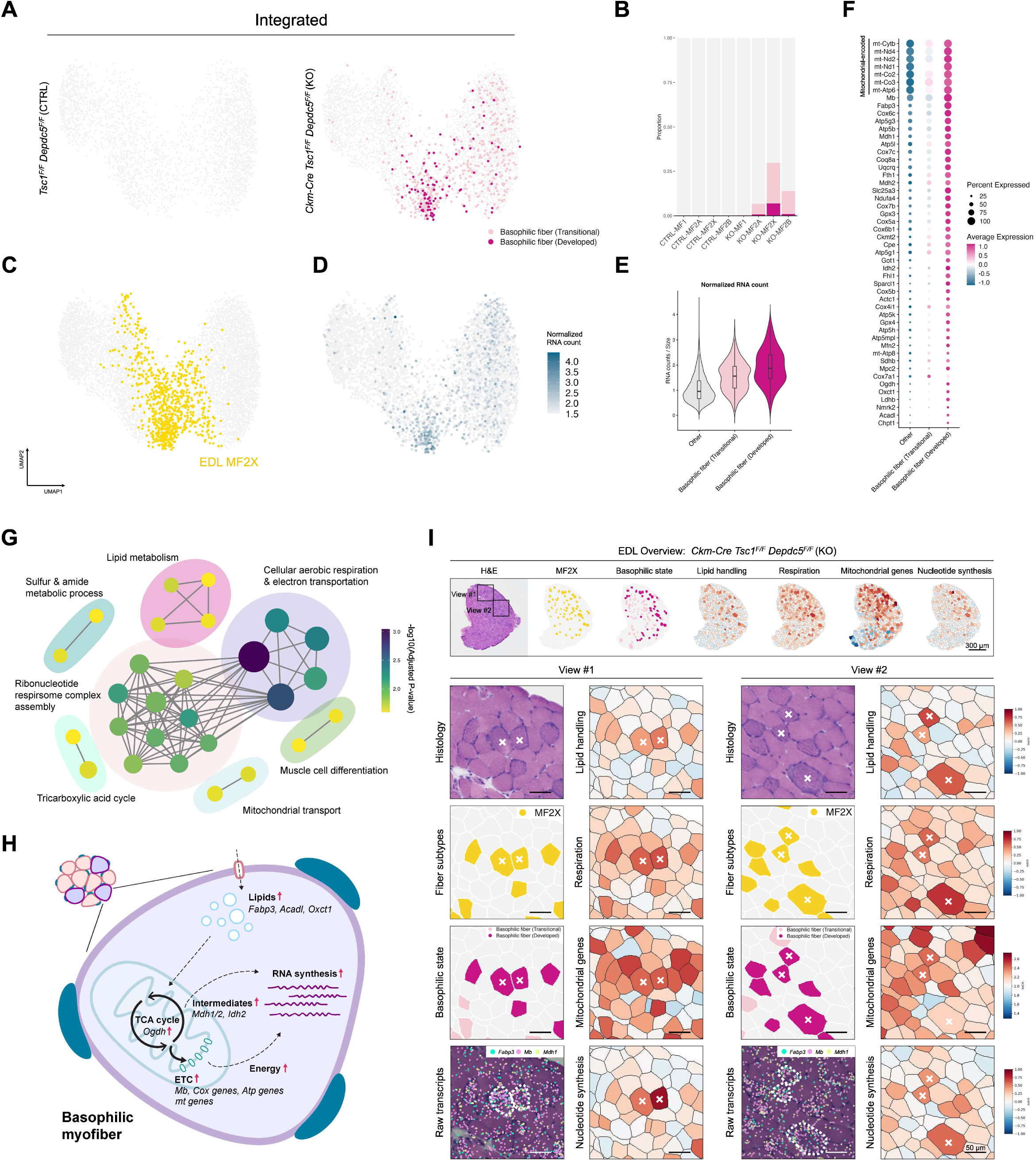
Basophilic fibers activate lipid and oxidative metabolism to sustain elevated RNA synthesis. (A) Integrated UMAP manifold, colored with transitional (pink) and developed (red) basophilic fibers, shown separately for control (left) and mutant (right) groups. (B) The prevalence of basophilia was quantified for each fiber type cluster. (C) UMAP manifold displaying MF2X myofibers originated from EDL muscles (yellow). (D) UMAP manifold displaying RNA density (size-normalized RNA count) of each myofiber. (E) Violin plot visualizing the distribution of size-normalized RNA counts for transitional (pink) and developed (red) basophilc fibers as well as other fibers (grey). (F) Dot plot visualizing the expression of genes upregulated in basophilic fibers. (G) GO enrichment network analysis based on genes specifically upregulated in basophilic fibers. Pathways were grouped by functional similarity. Node size corresponds to the number of genes associated with each pathway, and node color reflects enrichment significance (adjusted p-value). For the complete results with GO term names, see Fig. S5C. (H) Schematic diagram illustrating transcriptome regulation underlying mTORC1-induced basophilia in type IIx fibers in EDL. (I) Spatial images of a representative section from mutant EDL. Top: overview images presenting from left to right as H&E histology, MF2X identity, basophilic states, and cumulative score maps for following genes. Lipid handling score: *Fabp3, Acadl*, *Oxct1*, *Mfn2*, *Chpt1*, *Gpx4*, and *Ckmt2.* Respiration score: *Mb*, *Cox* genes, and *Atp* genes (see Fig. 5F). Mitochondrial gene score: mitochondrial genome-encoded genes (see Fig. 5F). Nucleotide synthesis score: *Idh2*, *Ogdh*, *Mdh1/Mdh2*, *Ldhb*, *Mpc2*, and *Nmrk2. Bottom:* representative basophilic fibers magnified in two views, corresponding to the boxes above in the H&E image. Raw transcripts are shown in colored dots as indicated in insets. Crosses and dotted borders highlight representative MF2X fibers. For the images from other mutant EDL sections, see Fig. S5A.

We next examined which genes were specifically expressed in basophilic fibers using differential expression analysis. Mitochondrial genome–encoded genes (Figure 5F; those beginning with ‘mt-’) were strongly upregulated, along with numerous additional transcripts. To systematically understand the molecular characteristics of basophilic fibers, we performed GO pathway enrichment analysis (Figure 5G). The result revealed a strong enrichment of genes mediating lipid metabolism, mitochondrial transport, tricarboxylic acid (TCA) cycle, electron transportation, and nucleotide synthesis (Figure 5G). These findings suggested a pronounced upregulation of a coordinated programs in basophilic fibers.

With further investigation, we identified multiple fatty acid and lipid metabolism-related genes, including *Fabp3* (fatty acid transport), *Acadl* (long-chain β-oxidation), *Oxct1* (ketone body catabolism), *Mfn2* (mitochondrial fusion and lipid exchange), *Chpt1* (phosphatidylcholine biosynthesis), and *Gpx4* (lipid peroxide detoxification), prominently upregulated in basophilic fibers (Figure 5F), reflecting a broad program enhancing lipid handling and mitochondrial lipid utilization. *Ckmt2* (mitochondrial creatine kinase) additionally linked lipid-derived energy flux to ATP buffering, bridging fatty acid oxidation with downstream energy production. Elevated expression of *Mb* (myoglobin), multiple *Cox* subunits (cytochrome c oxidase), *Atp* genes (ATP synthase), and several mitochondrially encoded respiratory chain components, as mentioned above, highlighted elevated mitochondrial respiration.

Enrichment of ribonucleotide metabolic pathways was also observed (Figure 5F, 5G, S5C). Several TCA-associated enzymes contributed both energy and building blocks for nucleotide production: *Idh2* (isocitrate dehydrogenase, generating NADPH for biosynthesis), *Ogdh* (α-ketoglutarate dehydrogenase, producing NADH while channeling carbon through the cycle), *Mdh1/Mdh2* (malate dehydrogenases, regenerating oxaloacetate that serves as a source of aspartate for nucleotide rings), and *Ldhb* (lactate dehydrogenase B, maintaining NAD⁺/NADH balance and controlling substrate flow into mitochondria). In parallel, *Mpc2* (mitochondrial pyruvate carrier) supported entry of glycolytic carbon into the cycle, while *Nmrk2* (nicotinamide riboside kinase 2) contributed to NAD⁺ salvage, sustaining redox reactions required for biosynthesis. Together, these signatures point to a coordinated program in which fatty-acid oxidation and respiration not only supply ATP but also drive TCA cycle flux that yields reducing power and carbon intermediates, thereby fueling purine nucleotide synthesis and ultimately resulting in the elevated RNA content of basophilic fibers (Figure 5H).

To illustrate this framework within a histological context, *Fabp3*, *Mb*, and *Mdh1* were highlighted as representative markers of lipid handling, oxidative respiration, and ribonucleotide metabolism, respectively (Figure 5I, raw transcripts view). In parallel, A selection of genes corresponding to these three pathways were combined into cumulative scores, and their relative expression levels were assessed and visualized (Figure 5I, S5A, right panel, S5D). This analysis confirmed the specific activation of all three pathways in basophilic fibers, which correspond to MF2X in EDL muscle. Taken together, our results indicate that mTORC1 hyperactivation reprograms MF2X fibers in EDL muscle toward lipid-supported respiration and nucleotide synthesis, giving rise to the basophilic phenotype.

### Stress-induced reprogramming of type IIb fibers in EDL reveals transcriptional adaptability

Type IIb fibers (MF2B), typically defined as the most glycolytic and fast-contracting population, have recently been shown to exhibit transcriptional variation under different contexts (19, 32). With deeper investigation of raw transcriptome data, we identified an internal heterogeneity among MF2B fibers that was masked by an aggressive batch correction applied during integration. To uncover this subtle yet critical diversity, we isolated and sub-clustered 4,182 myofibers from EDL, a muscle predominantly composed of MF2B fibers, without applying any batch correction (Figure S6A-E). Control EDL samples displayed minimal batch variation, whereas mutant samples showed somewhat greater variability across biological replicates, likely reflecting individual-specific responses or differences in mTORC1 hyperactivation-induced injury stages. Nonetheless, two mutant batches (KO-M1 & KO-M3) integrated well even without any batch correction, which could represent a reproducible transcriptome heterogeneity of MF2B myofibers (Figure S6C-D).

From this dataset, we identified a clear mTORC1 hyperactivation–driven subdivision of MF2B into three transcriptionally distinct groups among the two EDL replicates (Figure 6A-B). All three MF2B subgroups in mutant EDL exhibited elevated RNA content compared with controls, indicating active transcriptional responses (Figure S6F). Gene profiling revealed divergent programs: MF2B subtype-1 (58% of mutant MF2B) expressed stress-response transcripts resembling the global mTORC1 hyperactivation signature; MF2B subtype-2 (22%) largely escaped the stress response and instead showed increased expression of genes controlling mitochondrial and oxidative metabolism; and MF2B subtype-3 (20%) was characterized by re-expression of developmental and remodeling genes, including *Actc1* and *Myl4* (Figure 6C).

**Figure 6.**
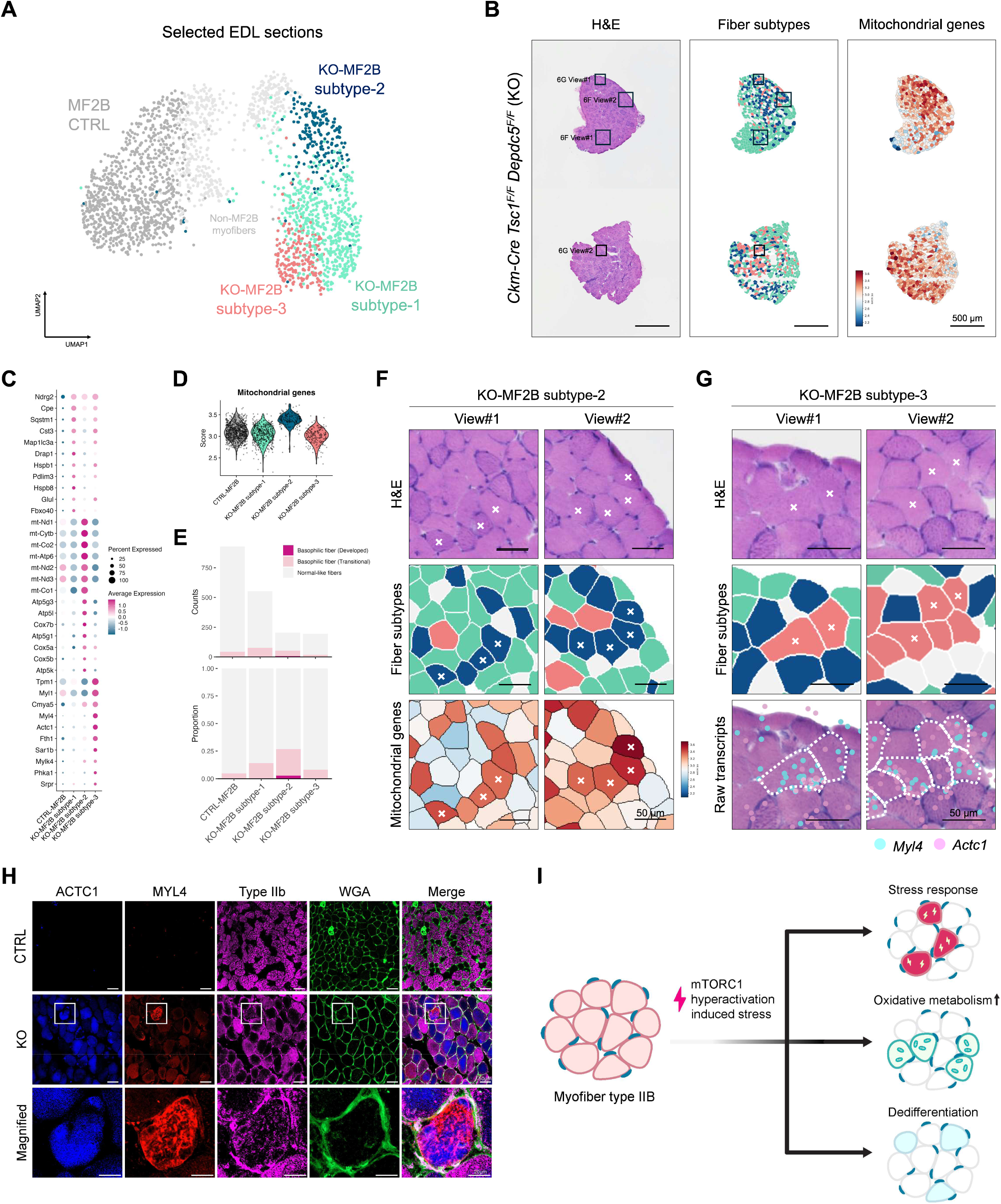
Type IIb fibers in EDL exhibit three distinct stress adaptation states without gross morphological alterations. Myofibers from EDL muscles were subset and analyzed to reveal MF2B heterogeneity. (A) UMAP manifold showed MF2B clusters in selected control and mutant EDL sections (n = 2). Control MF2B cluster and non-MF2B clusters are labeled in dark gray and light gray, respectively. For full analysis, see Fig. S6. (B) Spatial images of selected mutant EDL sections. Left: H&E histology. Middle: MF2B fiber subtype map, according to the clustering presented in (A). Right: Spatial map of a cumulative score created with the expression level of mitochondrial genome-encoded genes. Boxed areas in H&E images are magnified in Fig. 6F & 6G. (C) Expression of genes upregulated in each KO-MF2B subtype (1–3). (D) mitochondrial gene cumulative score compared between control and mutant MF2B subtypes. (E) Frequency of basophilic fibers in control and mutant MF2B subtypes. (F) Representative views of MF2B subtype-2 with histology (H&E), fiber-subtype map, and mitochondrial gene cumulative score map. Crosses indicate representative subtype-2 myofibers. (G) Representative views of MF2B subtype-3 with histology (H&E), fiber-subype map, and raw transcript plots of indicated genes. Crosses and dotted borders highlight representative subtype-3 myofibers. (H) Immunostaining of indicated proteins in control and mutant EDL muscles. WGA staining (green) delineates myofiber membranes. Boxed areas are magnified at the bottom. (I) Schematic diagram illustrating differential adaptation responses of MF2B fibers to mTORC1 hyperactivation-induced stress.

Among these subtypes, we focused on MF2B subtype-2, which suppressed stress response and rather upregulated oxidative metabolism. To quantitatively evaluate the mitochondrial content and activity, we generated a composite mitochondrial score based on all detectable mitochondrially encoded *mt* genes. The result confirmed that MF2B subtype-2 exhibited a relatively high mitochondrial level that is comparable to basophilic MF2X fibers, the oxidative population in EDL muscles (Figure 6C-D, S6H). However, interestingly, unlike basophilic MF2X fibers, MF2B subtype-2 fibers largely maintained normal morphology (Figure 6E–F). These findings suggest that a fraction of MF2B adopts oxidative metabolism without altering its myosin isoforms or developing pathological morphology in response to mTORC1 hyperactivation.

MF2B subtype-3, highlighted by developmental and de-differentiation signatures, did not display any obvious morphological phenotypes (Figure 6E & 6G). Transcriptional spatial mapping and quantitative analyses validated the elevated expression of *Actc1* and *Myl4* in this subgroup (Figure 6G, right column, S6I). Co-immunostaining of ACTC1 and MYL4 revealed both overlapping and non-overlapping fibers (Figure 6H), pointing to a more complex downstream regulatory mechanism.

Despite these wide variations in transcriptome differences between MF2B subtypes, they maintained their MF2B identity by robustly expressing its characteristic *Myh4* isoform (Figure S6B); *Myh4* expression level was very high across all MF2B subtypes without any notable differences between the subtypes.

Together, these findings demonstrate that type IIb fibers diverge into distinct transcriptional trajectories under mTORC1 hyperactivation (Figure 6I) while maintaining their fiber-type identity, highlighting previously unrecognized heterogeneity and transcriptional adaptability within a fiber type that was traditionally considered uniform.

### mTORC1 hyperactivation drives macrophage–fibroblast accumulation in soleus muscle

Aside from myofibers, or myocytes, many non-myocytic cell types—including fibroblasts, endothelial cells, and immune cells—reside within skeletal muscle, where they provide essential support for tissue maintenance and homeostasis. To investigate how these populations respond to mTORC1 hyperactivation, we profiled non-myocytes using 24-µm diameter (flat-to-flat width) hexagon spatial gridding. This revealed diverse cell types, including macrophages, fibroblasts, neuromuscular junctions (NMJs), and erythrocytes (RBCs) (Figure 7A-B).

**Figure 7.**
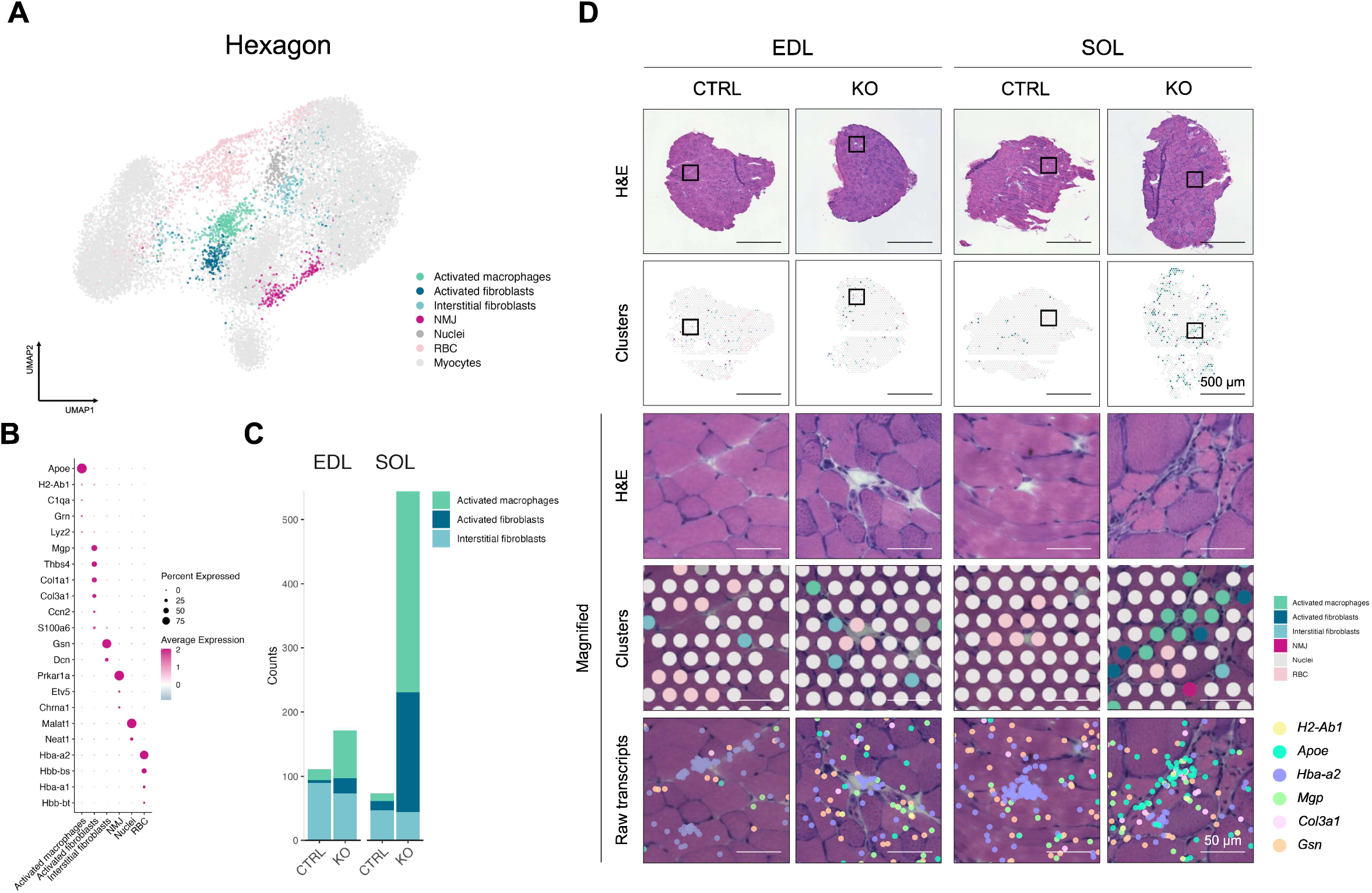
mTORC1 hyperactivation drives macrophage-fibroblast accumulation in SOL muscle. Non-myocytes were identified in skeletal muscle using 24 μm non-overlapping hexagon gridding. (A) UMAP manifold of the hexagonal grid dataset displaying non-myocyte clusters, conducted without batch correction. (B) Expression of non-myocyte markers in different cell types. (C) Quantification of macrophage and fibroblast cell type prevalence in control and mutant mice. (D) Spatial images showing H&E histology and cluster maps in representative muscle sections from each group. Boxed areas are magnified. Transcript expressions are displayed as colored dots as indicated.

While resting non-myocytic cell types are equally presented in all batches (Figure S7A-S7B), activated macrophages and fibroblasts were highly and specifically enriched in mutant SOL muscle (Figure 7C, S7C). Macrophage cluster strongly expressed *Apoe*, marking them as tissue-resident macrophages (TRMs), whereas activated fibroblasts, separated from *Gsn*-expressing interstitial fibroblasts, expressed *Mgp*, *Thbs4*, *Col1a1*, and *Col3a1*, along with relatively low *Gsn* expression (Figure 7B, S7D). These observations indicated the development of a fibrotic environment in the mTORC1-hyperactivated mutant SOL muscles, which are enriched with histopathological features such as myofiber hypertrophy and central nucleus. Importantly, activated TRMs were often found in close proximity to activated fibroblasts (Figure 7D), a spatial relationship that aligns with the established role of macrophages in driving fibrotic programs through paracrine signals that stimulate fibroblast activation and matrix deposition (39–41).

Overall, these results demonstrate that mTORC1 hyperactivation in myofibers not only drives intrinsic fiber pathology but also elicits secondary responses in non-myocytes within a specific muscle. The recruitment and activation of these cells is speculated to be triggered by altered tissue architecture and hypertrophic myofiber secretory signaling, creating a complex and interdependent pathological microenvironment.

## Discussion

Histopathological features such as basophilic fibers, fiber enlargement, and central nuclei have long served as hallmarks of skeletal muscle disease (38,42,43), yet the molecular programs underlying these morphologies remain poorly understood. Traditional approaches have struggled to connect phenotype with transcriptional state at the level of single fibers, while scRNA and snRNA sequencing lose their associated histological information (15–26). By combining high-resolution spatial transcriptomics with image-guided fiber segmentation, we establish a framework to directly link morphological phenotypes with their transcriptomes. This pathology-guided, single-fiber analysis reveals how sustained mTORC1 hyperactivation drives diverse and fiber-type–specific pathological responses across distinct muscle groups.

Even though the difference in susceptibility between EDL and SOL has long been recognized (44–46), our study is among the first to show that pathology-induced transcriptomic outcomes diverge sharply between these two muscles. In SOL, MF2X fibers predominantly underwent abnormal enlargement, whereas in EDL, MF2X fibers developed a basophilic phenotype. These contrasting trajectories indicate that MF2X fibers do not adopt a uniform pathological program but instead respond in a muscle-dependent manner. Given the distinct baseline fiber-type compositions and metabolic environments of SOL and EDL, it is plausible that local tissue context modulates how MF2X fibers respond to mTORC1 hyperactivation. In particular, the continuous activation and loading of SOL may contribute to its growth phenotype, whereas the relatively low usage of EDL might favor alternative pathological outcomes. It is also possible that MF2X fibers arising during development contain latent differences in identity, even if their baseline transcriptomes appear highly similar, which could contribute to muscle-specific outcomes under stress. Overall, these findings highlight the context-dependent plasticity of MF2X fibers and suggest that muscle-specific factors shape their adaptive responses to perturbation.

Through gene enrichment analysis, abnormally enlarged fibers were identified to engage sustained growth signaling, disrupted autophagic homeostasis, and structural remodeling (Fig. 4H), which is in line with the result from a previous study using bulk tissues (8). These processes were also shown to drive physiological myofiber hypertrophy in the context of mechanical overload (47). However, under continuous mTORC1 activation, they may tip the balance away from normal growth toward pathological remodeling. Even though hyperactive mTORC1 drives ongoing anabolic activity, including increased protein synthesis and ribosomal biogenesis, it simultaneously suppresses autophagy, thereby compromising proteostasis. This impaired capacity to clear damaged proteins and organelles likely exacerbates the imbalance between growth and quality control. Alongside these defects, cytoskeletal remodeling signatures, including reorganization of actin filaments and contractile apparatus components, suggest structural adaptation aimed at accommodating increased fiber size. Together, these processes form a foundation for hypertrophic expansion, but one that is unstable, as the combined disruption of proteostasis and autophagy predisposes the fibers to dysfunction and degeneration.

In EDL, MF2X fibers predominantly developed the basophilic phenotype, marked by lipid-supported respiration fueling nucleotide synthesis and culminating in elevated RNA content (Fig. 5H). Even though previous study has marked basophilia as a regenerative and metabolically oxidative phenotype (38), why these fibers adopt oxidative features in a glycolytic-dominant muscle under mTORC1 hyperactivation remains an open question. One possibility is that the oxidative shift arises as a means of redox management: by enhancing oxidative phosphorylation, fibers increase NADH and FADH₂ turnover, thereby alleviating oxidative stress. This process inevitably generates TCA intermediates that could, in principle, be directed into multiple metabolic fates. However, under mTORC1 hyperactivation, where anabolic programs strongly promote RNA and protein synthesis, these intermediates are likely funneled preferentially toward nucleotide production. Alternatively, the oxidative shift may serve a more directed purpose with fibers deliberately engaging lipid-supported respiration as the most efficient pathway to sustain nucleotide and protein synthesis. In either case, the phenomenon underscores the metabolic plasticity of MF2X fibers, which appear capable of recruiting oxidative programs to meet the demands of pathological growth within an otherwise glycolytic muscle environment.

Beyond basophilic and enlarged fibers, we also assessed additional pathological features commonly associated with myopathy. Fibers with central nuclei were more frequent in mutant muscle, consistent with regenerative remodeling (42), but our transcriptomic analysis did not yield a broad set of differentially expressed gene signatures for this nuclei population, likely reflecting technical difficulties or biological heterogeneity. However, we identified a small group of genes, including several zinc finger proteins, that were selectively enriched in central-nucleus fibers, suggesting potential roles in mediating central-nuclei–associated muscle pathology. We also noted that different muscle tissues display variability in phenotype prevalence, which we attribute to differences in the stage of mTORC1 hyperactivation across muscles. While all the sections were included in the integrative analysis, a subset of tissues were more extensively focused on to reveal fine-level heterogeneity that was masked during aggressive batch correction procedures.

In addition to these pathological features, our analysis also uncovered unexpected adaptability within MF2B fibers (Fig. 6I). Traditionally viewed as the most glycolytic and least flexible fiber type (48,49), MF2B in EDL diverged into three distinct subgroups under mTORC1 hyperactivation. One subgroup displayed the canonical stress-response program, consistent with the global transcriptomic changes induced by mTORC1 activation. The second subgroup, however, showed elevated expression of mitochondrial and oxidative genes while retaining a morphologically normal appearance. We speculate that this reflects a compensatory shift, in which a fraction of MF2B fibers adopt oxidative features to support aerobic metabolism that would normally be contributed by MF2X fibers, which are diverted into the basophilic trajectory. The third subgroup re-expressed developmental and structural regulators such as *Actc1* and *Myl4*, pointing to a remodeling program that may reflect partial de-differentiation or activation of regenerative pathways. These divergent outcomes suggest that mTORC1 hyperactivation does not impose a uniform response in glycolytic fibers, but instead interacts with local or intrinsic cues to channel MF2B into distinct transcriptional trajectories. With responses largely independent from overt morphological pathologies, we speculate that these heterogeneous programs may be shaped and affected by surrounding non-myocytes or neighboring fibers. Importantly, despite these wide variations in transcriptome response, they maintained their MF2B identity by expressing their characteristic myosin isoform *Myh4* at robust levels. These findings reveal that MF2B fibers, long considered metabolically rigid, can mount heterogeneous transcriptional responses when challenged, suggesting a previously underappreciated capacity for remodeling within this fiber type.

In conclusion, our study provides a comprehensive view of how sustained mTORC1 hyperactivation reshapes skeletal muscle at single-fiber resolution. By integrating pathology-guided segmentation with spatial transcriptomics, we uncovered fiber-type–specific adaptations, including divergent MF2X trajectories and unexpected heterogeneity within MF2B, as well as non-myocytic responses that promote fibrotic remodeling. These findings extend our understanding of the molecular programs that underlie classic histopathological features of myopathy and emphasize the plasticity of fiber types under stress. A limitation of our study is that it was performed in murine models of mTORC1 hyperactivation, and further work will be needed to determine whether similar adaptive programs operate in other pathological contexts or human diseases. Nonetheless, this work highlights the value of spatially resolved transcriptomics for linking morphology to molecular state and establishes a framework for future investigations into the mechanisms that govern muscle pathology.

## Data Availability

All source data, source codes for custom analyses in the paper and some processed datasets were deposited to Deep Blue repository (https://doi.org/10.7302/p25z-mk61).

## Methods

### Sex as a biological variable

In this study, spatial transcriptomic analyses were performed in male mice. Our previous characterization of the same mouse model showed that mTORC1-driven myopathy and associated cardiac and respiratory defects occur in both male and female mice (6,14), indicating that the core pathological consequences of mTORC1 hyperactivation are not sex restricted. Many functional and molecular assays in that work were carried out in males, and the present study builds on that dataset by adding high-resolution, fiber type-specific and muscle-specific transcriptional information in the same sex. Consequently, the spatial and single-myofiber programs described here are defined in male skeletal muscle, and it remains unknown whether the detailed histopathological and transcriptional trajectories are identical in females. Future studies that include both sexes and are prospectively powered for sex-stratified analyses will be required to determine whether sex modifies susceptibility to mTORC1 hyperactivation, the responses of individual fiber types, or associated non-myocytic remodeling.

### Rodent muscular mTORC1 hyperactivation model

As previously described (6,14), *Tsc1^F/F^*and *Ckm-Cre* mice (C57BL/6J background) were obtained from Jackson Laboratories (stock no. 005680), and *Depdc5^F/F^* mice were collected from the European Mouse Mutant Archive (EM:10459). *Tsc1^F/F^ Depdc5^F/F^* double knockout mice were generated by interbreeding, and progeny were subsequently crossed with *Ckm-Cre* mice to produce skeletal muscle–specific knockout animals. All mice were maintained under protocols approved by the University of Michigan Institutional Animal Care and Use Committee (IACUC). To minimize batch effects, four control and four knockout littermates were selected for tissue harvest. Immediately after dissection, muscle tissues were embedded in O.C.T. compound (23-730-571, Fisher Scientific) and snap-frozen in liquid nitrogen–cooled 2-methylbutane (MX0760, Sigma-Aldrich).

### Seq-Scope array production (1st-Seq)

Seq-Scope procedures were described in detail previously (30,31). Seq-Scope is a solid-phase transcriptome spatial capture system that has undergone several upgrades. The latest version was built upon Illumina NovaSeq 6000 platform with a 7 mm × 7 mm imaging area. Barcoded capture probe clusters on the array surface were generated using HDMI32-DraI, a custom single-stranded oligonucleotide library (IDT), and Read1-DraI sequencing primer through a sequence-by-synthesis strategy. The sequencing output FASTQ file generated during array construction contained the barcode sequences and their corresponding XY coordinates.

The array originated from Illumina NovaSeq 6000 flow cell contains four channels with 7 mm × 70 mm imaging areas. Arrays were split and diced into suitable sizes for storage and experimental use. Before application, arrays required several preprocessing steps. Arrays were first washed three times with nuclease-free water, followed by overnight incubation at 37 °C with a DraI (R0129, NEB) / CIAP (M0525, NEB) enzyme mixture. The following day, arrays were treated with Exonuclease I (M2903, NEB) for 45 min at 37 °C, then washed sequentially: three times with nuclease-free water, three times with 0.1 N NaOH for 5 min each, and three times with 0.1 M Tris (pH 7.5).

### Seq-Scope spatial transcriptome library generation and sequencing (2nd-Seq)

Seq-Scope library generation was performed according to our previous paper (30). OCT-mounted frozen muscle blocks stored at –80 °C were equilibrated in a cryostat (Leica CM3050S) at –15 °C for 1 h prior to sectioning. Sections were cut at 10 μm thickness with a 5° cutting angle and placed onto cold Seq-Scope arrays, then warmed to room temperature to promote tight attachment. Tissue fixation was performed on the array with 4% formaldehyde (15170, Electron Microscopy Sciences) at room temperature for 10 min, followed by three PBS washes. Sections were then co-stained with DAPI (Invitrogen, D21490) and Alexa Fluor-488 WGA (Invitrogen, W11261) for 15 min at room temperature, washed three times with PBS, mounted in 5% glycerol, and imaged on a Keyence digital darkroom system for whole-tissue DAPI/WGA capture. The coverslip was then gently removed to avoid tissue damage, after which hematoxylin and eosin (H&E) staining was performed and whole-tissue H&E images were captured with a Keyence microscope.

Tissue permeabilization was carried out by incubating sections with 0.2 U/μL collagenase I (17018-029, Thermo Fisher) at 37 °C for 20 min, followed by 1 mg/mL pepsin (P7000, Sigma) in 0.1 M HCl at 37 °C for 10 min. Reverse transcription was performed between the permeabilized tissue and array surface using an RT mixture [1× RT buffer (EP0751, Thermo Fisher), 4% Ficoll PM-400 (F4375-10G, Sigma), 1 mM dNTPs (N0477L, NEB), RNase inhibitor (30281, Lucigen), Maxima H-RTase (EP0751, Thermo Fisher)] and incubated overnight at 42 °C in a humidified chamber. The following day, unbound single-stranded DNA probes were digested with Exonuclease I (M2903, NEB). Tissue was removed by incubation with a digestion cocktail (100 mM Tris, pH 8.0; 100 mM NaCl; 2% SDS; 5 mM EDTA; 16 U/mL Proteinase K, P8107S, NEB) at 37 °C for 40 min. Arrays were then washed three times each with nuclease-free water, 0.1 N NaOH (5 min per wash), and 0.1 M Tris, pH 7.5.

Library generation was performed immediately after washing. Arrays were incubated for 2 h at 37 °C in a humidified chamber with a library synthesis mixture [1× NEBuffer-2 (NEB), 10 μM TruSeq Read2-conjugated random primer (IDT), 1 mM dNTPs (N0477, NEB), Klenow Fragment (M0212, NEB), nuclease-free water]. The arrays were then washed with nuclease-free water, and libraries were collected twice by 0.1 N NaOH elution (5 min each). Eluates were neutralized with 3 M potassium acetate, pH 5.5, and purified with AMPure XP beads (1.2× bead/sample ratio, A63881, Beckman Coulter) according to the manufacturer’s instructions.

Libraries were amplified in two rounds of PCR using Kapa HiFi HotStart ReadyMix (KK2602, KAPA Biosystems). The first round used 2 μM TruSeq forward and reverse primers as described previously (30,31), and products were purified with AMPure XP beads (1× bead/sample ratio). In the second round, TruSeq indexing primers were applied, and products were size-selected with AMPure XP beads (0.6× bead/sample ratio). Library quality was assessed on an Agilent 2100 Bioanalyzer (Agilent), with additional purification performed if necessary. Final libraries were sequenced using paired-end 100-cycle reads.

### Raw data processing

Seq-Scope data processing was using the NovaScope pipeline, as described in detail previously (30). 1st-Seq data table was generated from the 1st-Seq FASTQ files, containing array region lookup indices, quality-filtered barcode sequences, and corresponding spatial coordinates. The 2nd-Seq FASTQ reads were then processed against this table by (i) assigning array indices through region lookup, (ii) mapping reads to the spatial barcode map using spatial barcode sequences (HDMI), (iii) filtering barcoded reads with validated barcode sequences, and (iv) assigning spatial coordinates to the identified reads. Reads with assigned spatial information were aligned to the genome using STAR (50), and a spatial digital gene expression (sDGE) matrix was generated.

### Histology and sDGE alignment

H&E images were aligned to sDGE matrix using the *historef* package (version 0.1.3), which detects fiducial marks visible in both modalities to guide image registration, as described previously (30). DAPI/WGA images were subsequently aligned to the processed H&E images using Georeferencer function in QGIS (version 3.22.9).

### Myofiber segmentation and hexagon gridding

Myofiber segmentation was performed in Cellpose (51) using WGA-stained myofiber membranes as input. A custom segmentation model was trained to optimize fiber identification, and manual adjustments were applied when necessary. Segmentation masks generated by Cellpose were exported as NumPy arrays and processed with custom Python scripts to label and isolate individual fibers. Transcripts within each segmented myofiber were aggregated and stored as an aggregated spatial digital gene expression matrix. Longitudinal and oblique myofibers, which are minimally presented within the cross sections, were ruled out during pixel size counting.

For non-myocyte analysis, spatial hexagon grids (24 μm, non-overlapping) were generated using NovaScope, where transcripts within each grid were aggregated and treated as a single unit, with spatial coordinates assigned to the hexagon center.

### Data Analysis and Visualization

Single-myofiber datasets were analyzed using Seurat package (52,53). Myofibers with low feature counts (nFeature cutoff: 200) were removed to optimize clustering performance. Each batch was normalized with SCTransform, and datasets were integrated using SCT-normalized values. Principal components were calculated using the RunPCA function, and high-quality components were selected to generate UMAP embeddings, followed by cluster identification with the FindNeighbors and FindClusters functions. Myofiber clusters were annotated based on marker gene expression, and unique transcript signatures were identified with the FindMarkers and FindAllMarkers functions.

Gene ontology enrichment analysis was performed with ClusterProfiler (54) using the org.Mm.eg.db annotation (Carlson, M., Bioconductor). Data visualization was supported by various R packages including Seurat, ggplot2, ggvenn (Yan, G., version 0.1.10), ggVennDiagram (55), ComplexUpset (krassowski, M., version 1.3.3), ComplexHeatmap (56) and enrichplot (54). Spatial myofiber projections were generated from segmentation NumPy arrays using custom python scripts. Raw spatial gene expression was visualized with a custom software platform that displayed both histology and the sDGE matrix.

Hexagon-based spatial transcriptomes (24 μm non-overlapping grids) were analyzed independently without dataset integration, primarily to detect non-myocytes. Each hexagon was treated as a single cell. Normalization and clustering were performed in Seurat using SCTransform, RunPCA, and RunUMAP function, followed by marker-based annotation.

### Statistics

Genes significantly upregulated in mutant muscle comparing to control muscle were first filtered using stringent cutoffs on expression level and fold-change, followed by systematic classification on fiber types using a combination of Kruskal–Wallis and pairwise Wilcoxon tests. The Kruskal–Wallis test identified genes with significantly higher expression in their top-expressing fiber type, thereby defining fiber type–specific signatures (adjusted p.value < 0.05). The Pairwise Wilcoxon tests were subsequently conducted to compare expression levels across fiber types, allowing us to define partially shared gene groups as well as globally enriched transcripts (adjusted p.value < 0.05).

### Immunohistochemistry

Frozen muscle sections were equilibrated to room temperature before processing. Sections were permeabilized with 0.5% Triton X-100 in PBS for 5 min, followed by three PBS washes (5 min each). Fiber-type immunostaining was performed using the Mouse on Mouse (M.O.M.) Immunodetection Kit (VectorLabs, BMK-2202) according to the manufacturer’s instructions. Briefly, sections were incubated overnight at 4 °C with M.O.M. Mouse Ig Blocking Reagent, then with primary antibodies diluted in M.O.M. Diluent at optimized concentrations. Alexa Fluor–conjugated secondary antibodies (Invitrogen) were diluted in M.O.M. Diluent and applied for 60 min at room temperature. For membrane visualization, sections were incubated with Alexa Fluor 488–conjugated WGA (Invitrogen, W11261) for 10 min prior to mounting with ProLong Gold Antifade Mountant (Invitrogen, P36934). Images were acquired using a Zeiss LSM 980 confocal microscope equipped with an Airyscan 2 detector (Carl Zeiss Microscopy). Primary antibodies: BA-D5 (MF1), SC-71 (MF2A), BF-F3 (MF2B) were acquired from Developmental Studies Hybridoma Bank (DSHB), University of Iowa. ACTC1 (66125-1-IG) and MYL4 (67533-1-IG) were purchased from ProteinTech. UCHL1 (PA5-29012) and IGFBP5 (PA5-37369) were purchased from Invitrogen. SQSTM1/p62 (5114) were purchased from Cell Signaling.

---

**Figure S1.**
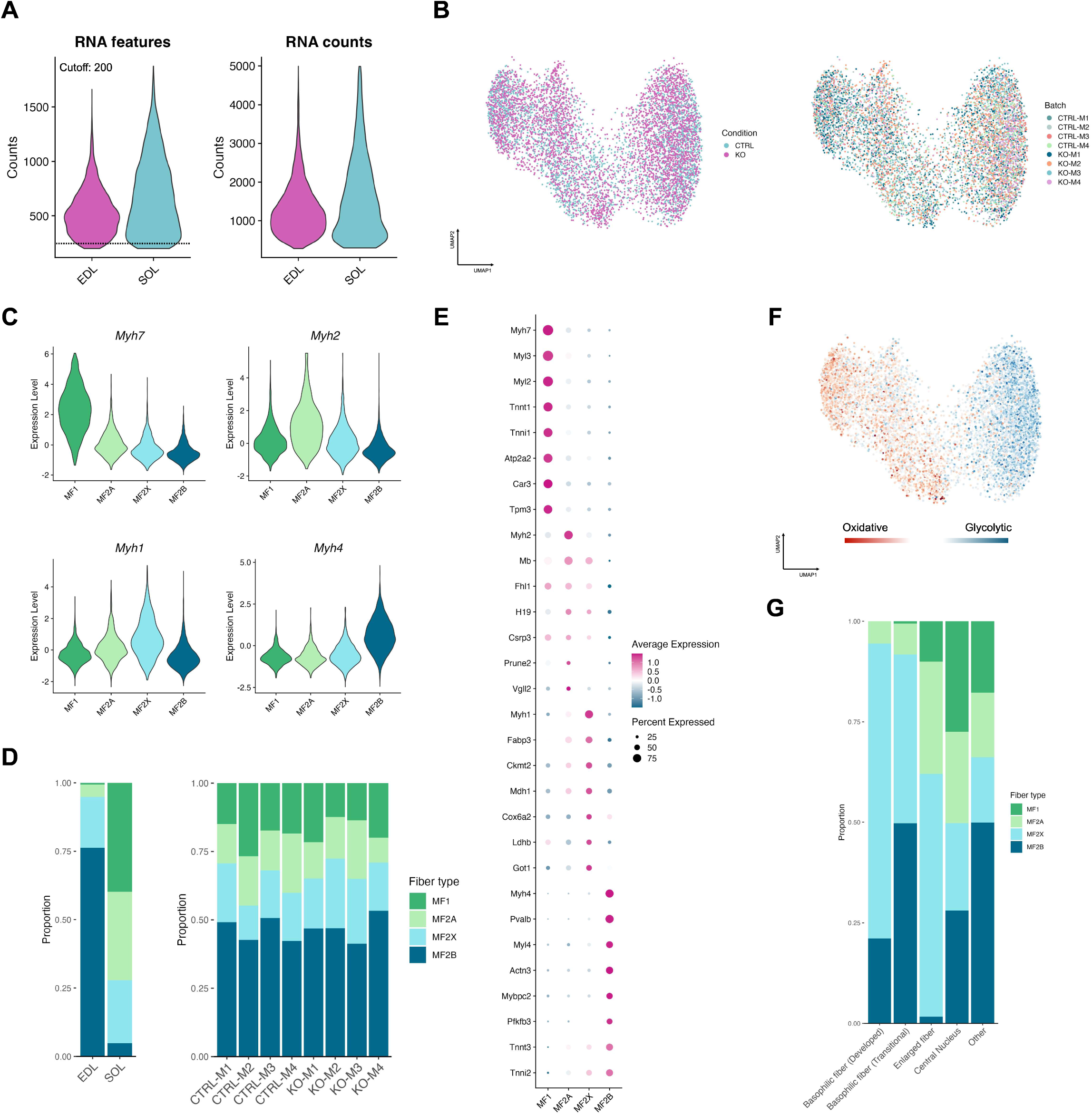
Stratification of single-myofiber transcriptome into classical myofiber types. (A) Distribution of unique RNA features (number of detected genes; left) and transcript counts (right) per segmented myofiber, shown separately for EDL and SOL. A minimum cutoff of 200 unique RNA features was applied prior to integration and downstream analyses. (B) Data integration performance. The UMAP manifold is colored according to the conditions (left) and batches (right) of individual myofibers. All eight biological replicates, from either control or mutant mice, contributed to all clusters with similar proportions. (C) Expression of myofiber-specific myosin heavy chain isoforms across designated myofiber types. (D) Proportion of myofiber types across muscles (left) and batches (right). (E) Dot plot representing the expression of fiber type-specific marker genes in each myofiber type cluster. Dot size represents the proportion of myofibers expressing each gene, and dot color represents the average expression level. (F) UMAP manifold colored according to the combined scores of metabolic-related genes across the oxidative (red) and glycolytic (blue) axis. (G) Distribution of myofiber types within each pathological phenotype category.

**Figure S2.**
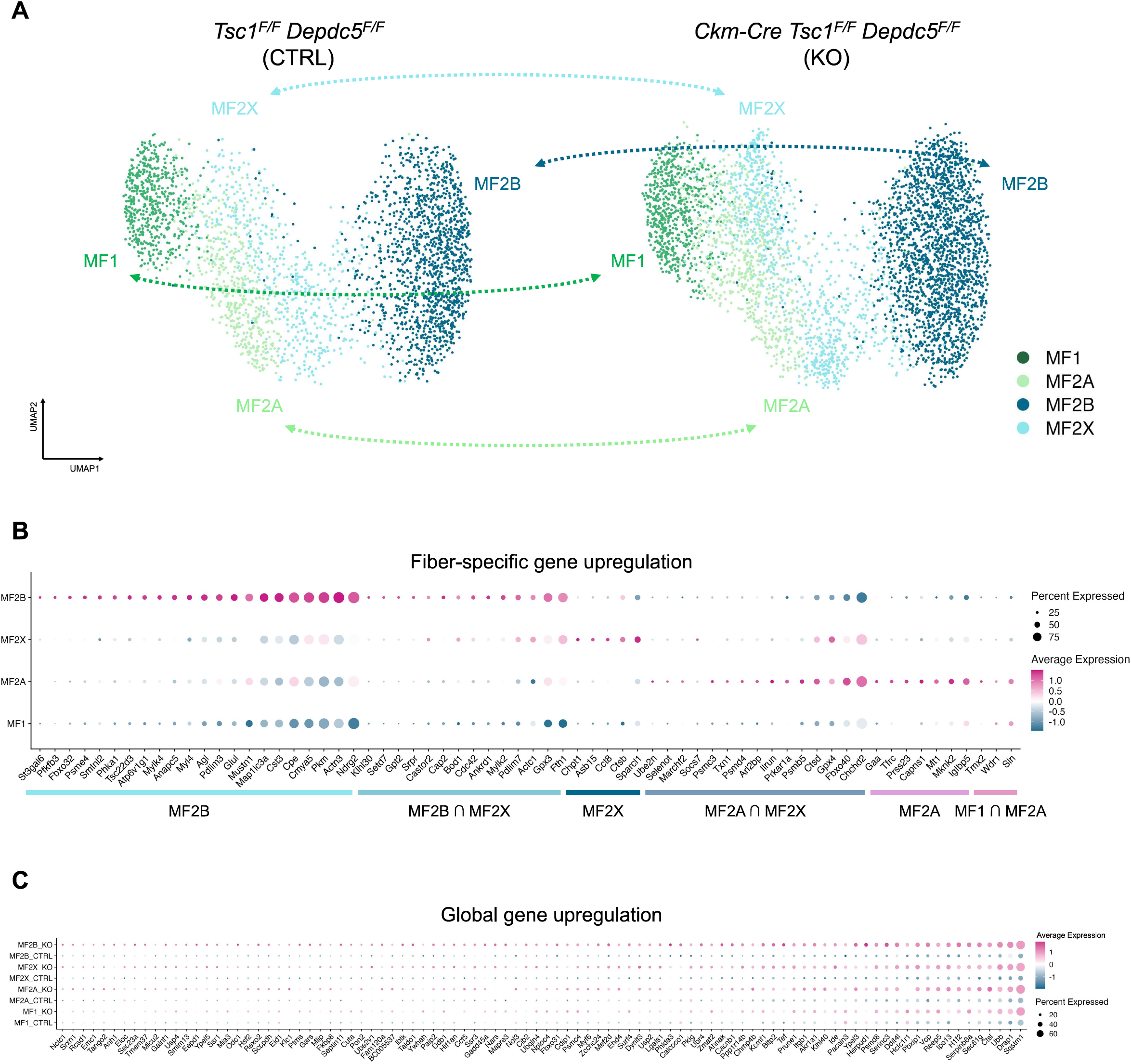
Validation of fiber type-specific and global gene expression comparisons. (A) From the UMAP manifold showing fiber type identities (Fig. 1C), two condition-specific manifolds were generated. Dotted double-arrow lines indicate the fiber type–matched comparisons used for differential expression analysis. (B) Expression of mTORC1 hyperactivation-induced genes that are upregulated only in specific myofiber types in mutant muscle. The intersection symbol (∩) indicates genes that are upregulated in both of the indicated myofiber types. (C) Expression of globally upregulated genes induced by mTORC1 hyperactivation across all myofiber types.

**Figure S3.**
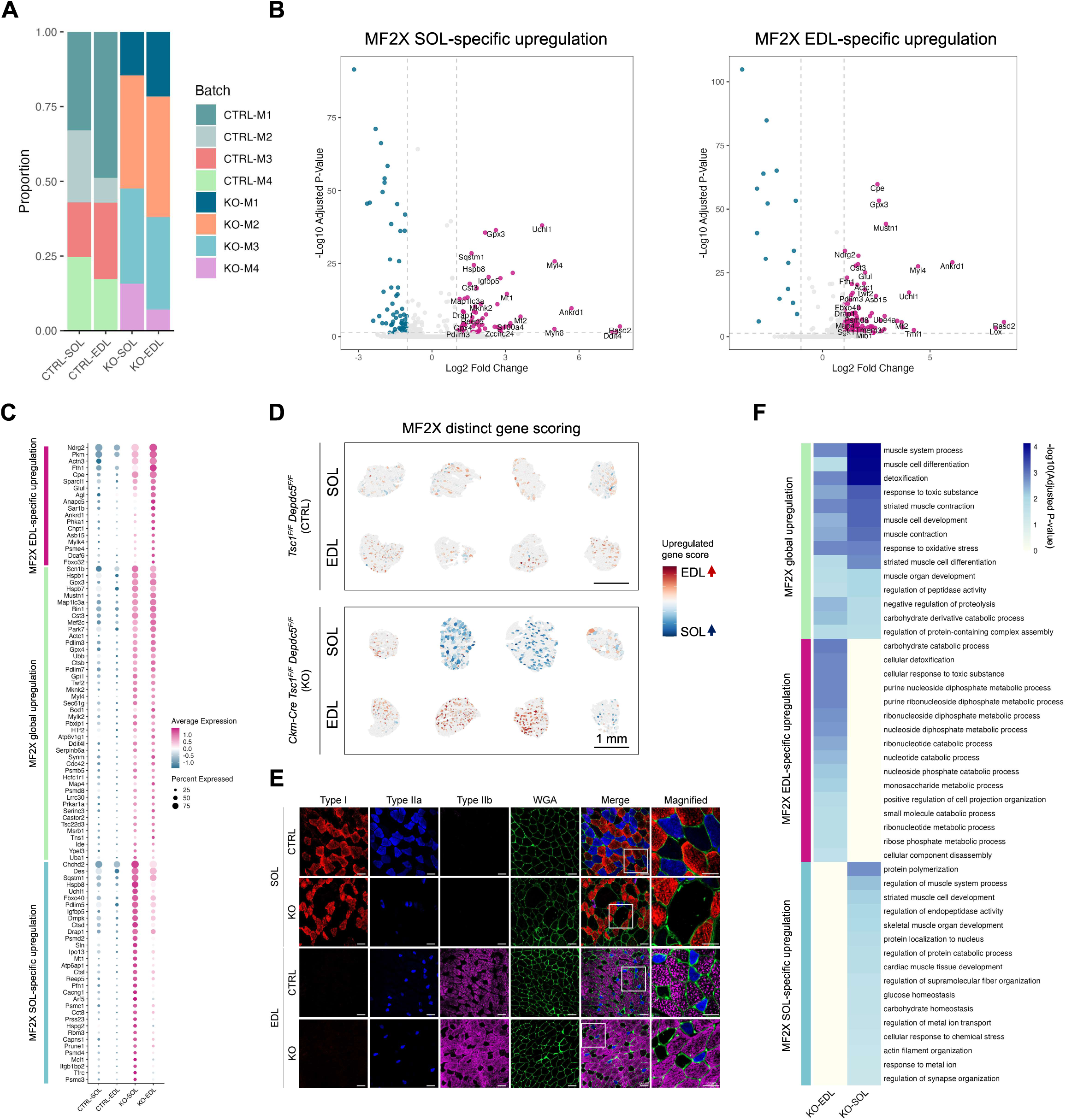
Validation of MF2X fiber responses under mTORC1 hyperactivation. (A) Contribution of different experimental batches to each indicated dataset of MF2X myofibers. (B) Differential expression analyses of MF2X fibers in SOL (left) or EDL (right) between control and mutant mice. Significantly upregulated or downregulated genes (adjusted p-values < 0.05, Log2FC > or < 1.5) are highlighted. (C) Expression of the complete list of genes that are upregulated in response to mTORC1 hyperactivation in all MF2X fibers (MF2X global upregulation) or specifically in MF2X of EDL or MF2X of SOL. Exclusivity was determined using pairwise Wilcoxon tests (adjusted p-value < 0.05). (D) Spatial images of muscle-specific MF2X cumulative module scores (MF2X distinct gene scoring) representing the axis of SOL-and EDL-specific responses. (E) Immunostaining of indicated fiber types in control or mutant EDL or SOL muscle. Type I (red), IIa (blue) and IIb (magent) fibers are visualized through specific antibody staining while type IIx fibers appear unstained (black). WGA staining (green) outlines myofiber membranes and extracellular matrix. Boxed areas in Merge column are magnified at right. (F) Heatmap showing the result of gene set enrichment analysis of genes upregulated in response to mTORC1 hyperactivation in all MF2X fibers (MF2X global upregulation) or specifically in MF2X of EDL or MF2X of SOL. Color represents the enrichment significance (adjusted p-values). The result was used to construct the GO enrichment network shown in Fig. 3H.

**Figure S4.**
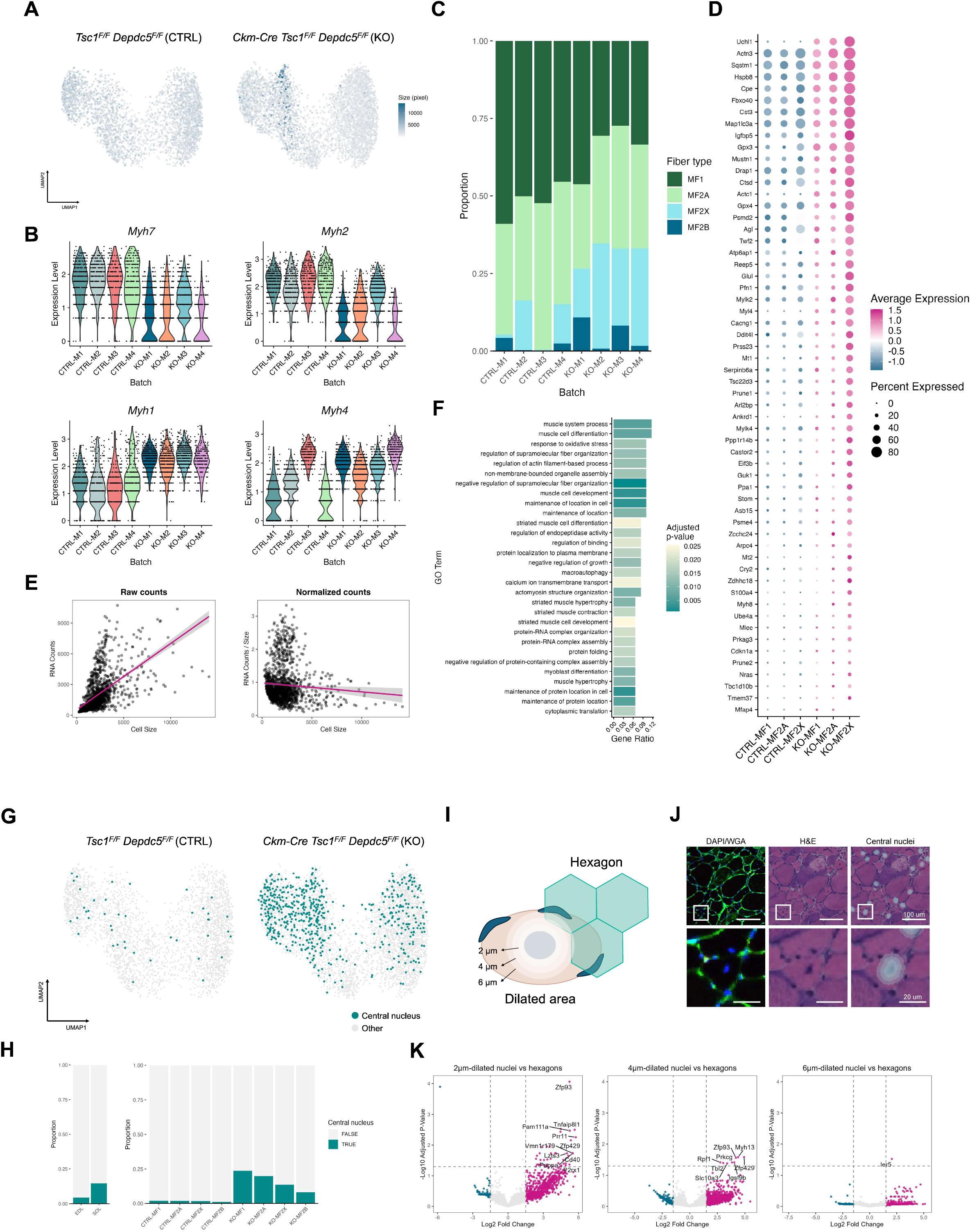
Transcriptomic characteristics of pathological features including myofiber enlargement and central nuclei. (A) Myofiber size visualized in integrated UMAP space, shown separately for control (left) and mutant (right) groups. (B) Expression of myofiber-specific myosin heavy chain isoforms across control and mutant SOL muscle batches. (C) Proportion of myofiber types in control and mutant SOL muscle batches. (D) Expression of mTORC1 hyperactivation-driven upregulated genes in SOL myofiber types. Most of these genes show highest expression in MF2X fibers in mutant mice. (E) RNA count-fiber size correlation. Right: correlation between transcript counts per myofiber (RNA Counts) and fiber size. Left: correlation between size-normalized transcript density (RNA Counts / myofiber) and fiber size (Cell size). (F) GO enrichment analysis of genes upregulated in enlarged fibers, corresponding to the GO network shown in Fig. 4G. (G) Centrally nucleated myofibers are marked in integrated UMAP space, separated by control (left) and mutant (right) groups. (H) Proportion of centrally nucleated myofibers in indicated subsets. (I) Histology-based spatial single-nuclei–like analysis. Central nuclei were identified from the histology images, and central nuclear transcriptome was extracted from spatial transcriptome with different radial ranges, reflecting lateral diffusion of transcripts. Analysis compared whole-tissue transcriptomes (segmented by 24 μm hexagon gridding) with central-nuclei transcriptomes. (J) Identification of centrally nucleated fibers by combining DAPI/WGA and H&E staining. Central nuclei with different radial ranges are visualized in a gradient on top of H&E images. Boxed areas are magnified below. (K) Comparison of central-nuclei transcriptomes with whole-tissue transcriptomes. Differentially expressed genes between tissue transcriptome and central nuclei transcriptomes of different radial ranges were shown in volcano plots. Upregulated (red) and downregulated genes (green) (Log_2_FC > 1.5 on either side) are highlighted; significantly upregulated genes (adjust p-value < 0.05) are labeled.

**Figure S5.**
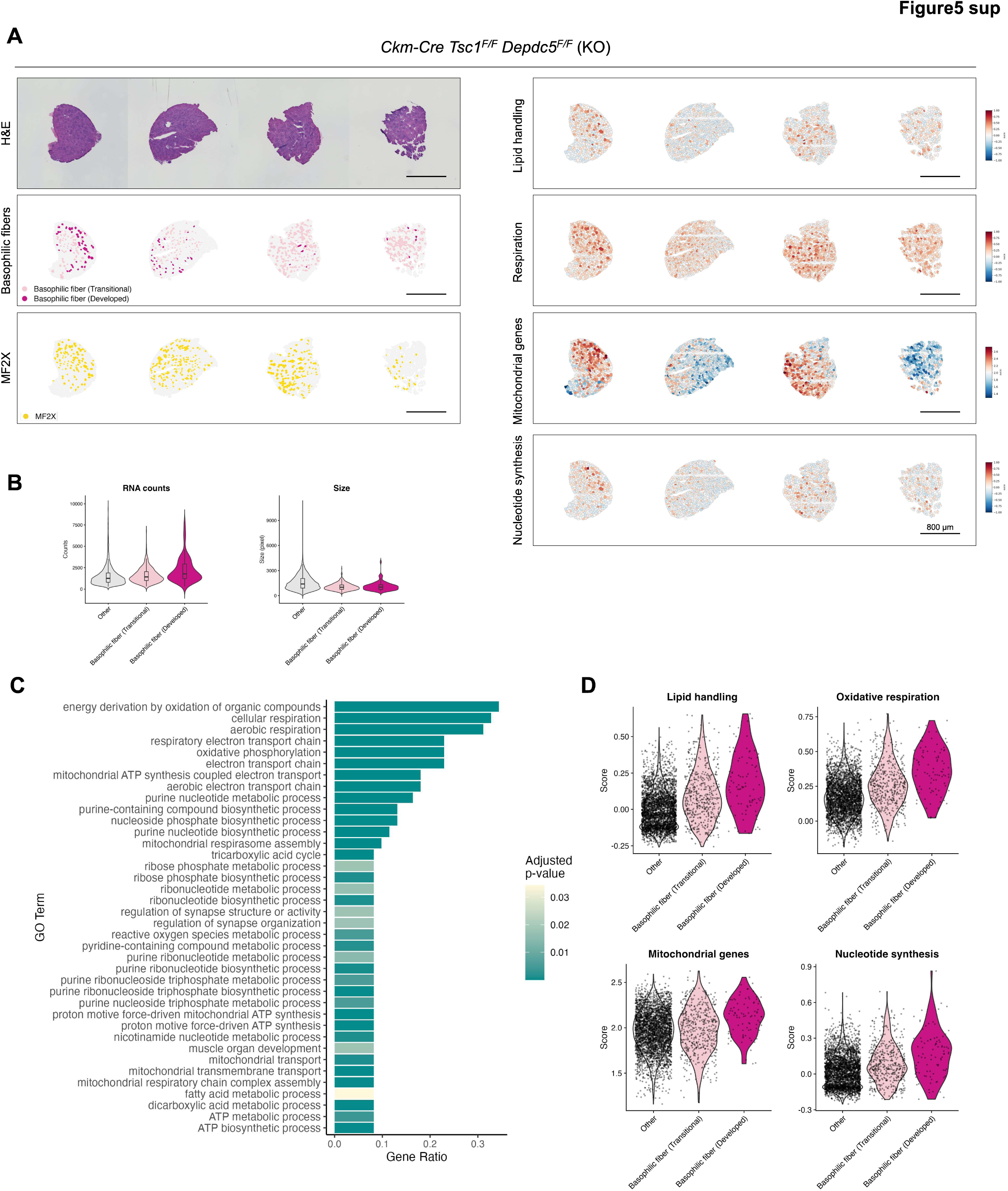
Basophilic fibers in EDL upregulate oxidative metabolism and RNA synthesis. (A) Spatial images of mutant EDL sections (full set). Left panel from top to bottom: H&E histology, basophilia map, and MF2X cluster map. Right panel from top to bottom: maps of lipid-handling module score, respiration module score, mitochondrial gene module score, and nucleotide synthesis module score, calculated as in Fig. 5I. (B) Distribution of RNA counts and fiber sizes across basophilic fiber states. (C) Enriched GO terms corresponding to nodes in the basophilic fiber functional network (Fig. 5G). (D) Distribution of cumulative module scores across basophilic fiber states.

**Figure S6.**
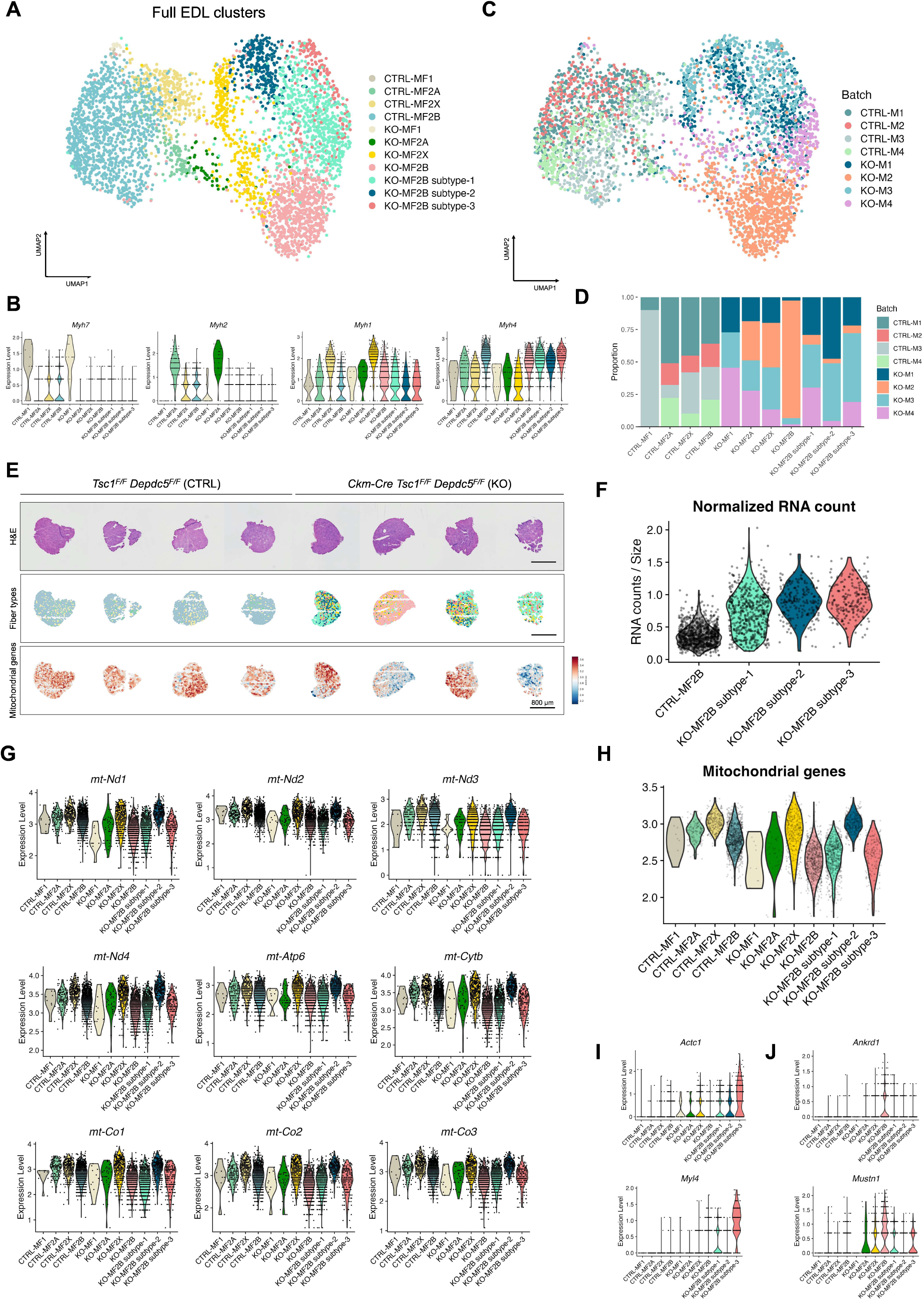
Diversification of MF2B responses to mTORC1 hyperactivation. (A) UMAP manifold displaying myofiber types from all EDL sections, constructed without batch correction. (B) Expression of myofiber-specific myosin heavy chain isoforms across designated myofiber types. All MF2B myofiber subtypes express Myh4, a key marker of MF2B. (C) UMAP manifold displaying experimental batches from all EDL sections, constructed without batch correction. (D) Proportion of individual batches in different myofiber types, presented in a bar graph. (E) Spatial images of 8 EDL sections (4 control and 4 mutant) used for the analyses in (A-D). H&E histology (top), fiber type map (middle), and mitochondrial gene cumulative score map (bottom) are shown. (F) Distribution of size-normalized RNA counts in different MF2B subclusters. (G) Expression of mitochondrial genome-encoded genes in different myofiber clusters of EDL. (H) Mitochondrial gene cumulative score in different myofiber clusters of EDL. (I-J) Expression of indicated genes in different myofiber clusters in EDL, representing the mutant MF2B subtype-3 (I) or the KO-M3 batch effect (J).

**Figure S7.**
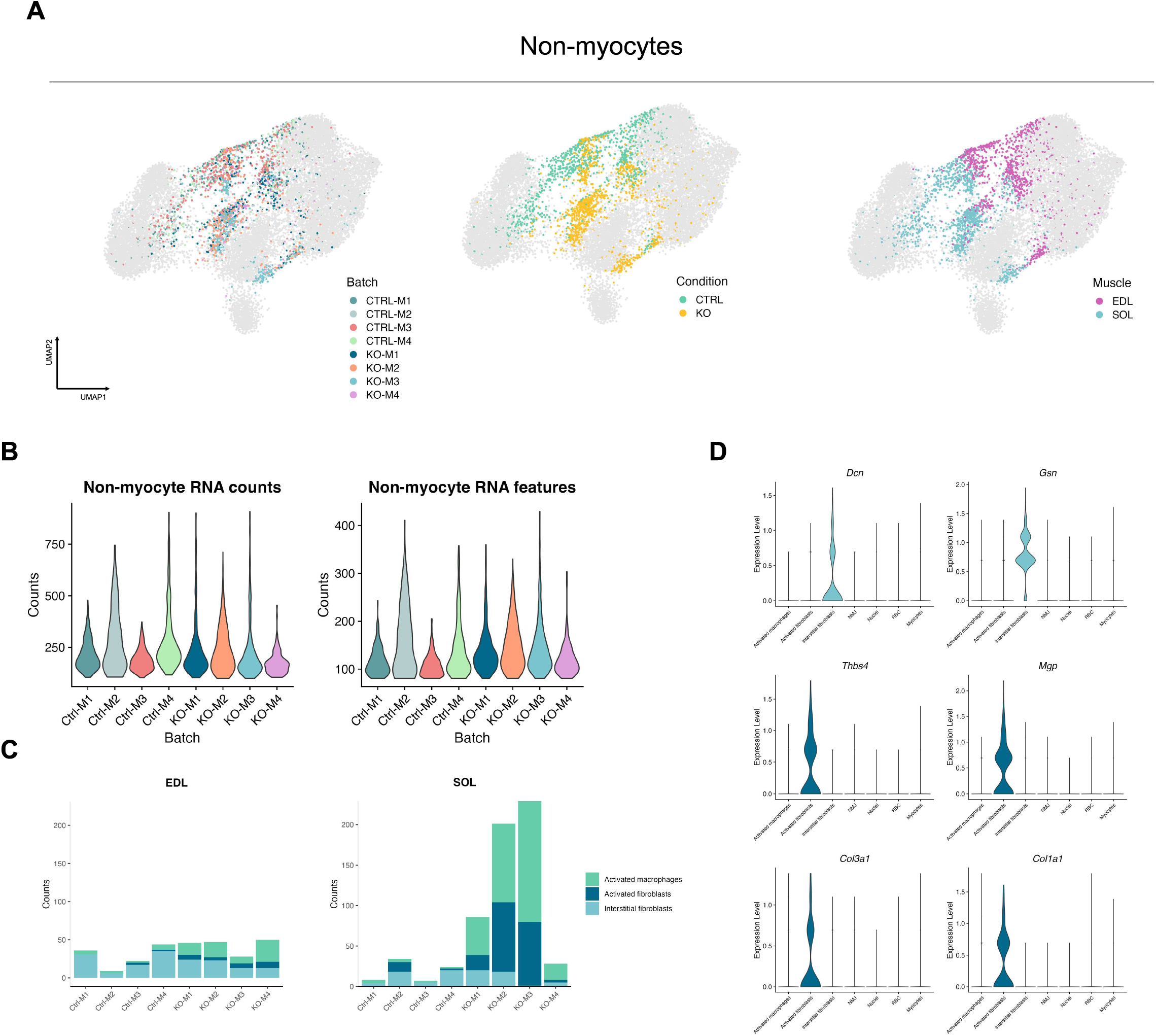
Hexagonal grid analyses of non-myocyte populations. (A) UMAP manifold, colored with batches (left), conditions (middle), or muscle (right), constructed without batch correction. Non-myocytes are highlighted in indicated colors, while myocytes are greyed out. (B) Distribution of unique transcript counts (left) and gene numbers (RNA features, left) per non-myocytes of indicated batches. (C) Quantification of macrophage and fibroblast cell type prevalence in indicated batches. (D) Expression of fibroblast activity-associated genes across non-myocyte clusters.

